# PHYCUT: Scalable multiplex CRISPR/Cas9 editing for genome engineering in the diatom *Phaeodactylum tricornutum*

**DOI:** 10.1101/2025.10.31.685893

**Authors:** Emily E. Stuckless, Lian S. Gai, Samuel S. Slattery, Kira H. Dempsey, Tyler S. Browne, Gregory B. Gloor, David R. Edgell

## Abstract

Diatoms are globally significant microalgae that contribute ∼ 20% of oxygen production and exhibit remarkable metabolic diversity. The marine diatom *Phaeodactylum tricornutum* has emerged as a promising synthetic biology platform for bioproduction of recombinant proteins, supported by a human-like *N*-linked glycosylation pathway. However, its *α*(1,3)-linked core fucose is immunogenic in humans and thus limits biopharmaceutical applications. One hurdle to efficient genome engineering in *P. tricornutum* is the lack of a robust system for simultaneous CRISPR/Cas9 editing at multiple sites. To overcome this limitation, we develop PHYCUT (***<u>Ph</u>****aeodactylum tricornutum* Cs**<u>y</u>**4-**<u>C</u>**as9 m**<u>u</u>**ltiplex **<u>t</u>**ool), a versatile plasmid-based CRISPR/Cas9 system that uses the Csy4 endoribonuclease to process multi-guide RNA arrays. To highlight PHYCUT applications, we demonstrate multiplex editing of all three *FucT* genes responsible for *α*(1,3) fucosylation in *P. tricornutum*, yielding strains with markedly reduced fucosylation of secreted proteins. PHYCUT enables facile, multiplexed genome engineering in diatoms and provides a foundation for humanizing the *P. tricornutum* glycosylation pathway to support next-generation algal biotechnology.

## INTRODUCTION

Diatoms are an ecologically important group of eukaryotic microalgae, estimated to be responsible for ∼ 20% of global oxygen productivity^1,2^. With their metabolic diversity and their ability to thrive and adapt to different environments, diatoms have garnered interest as biotechnological platforms with the potential to produce high-value compounds at low cost. *Phaeodactylum tricornutum* is a marine pennate diatom with a growing toolbox for genetic engineering, and the capacity to be grown in scalable photobioreactors. A fully-sequenced genome^3,4^, characterized promoter and terminator elements^5–7^, efficient DNA delivery protocols based on conjugation^8^, electroporation^9^, or biolistic transformation^10^, and CRISPR-Cas9 gene-editing tools^5,11–13^ empower *P. tricornutum* as a platform for biotechnology.

Biotechnological applications of *P. tricornutum* include pigment extractions for nutraceuticals and cosmetics^14^, strain-development for biofuel production^15,16^ and the production of recombinant proteins for diagnostics or biopharmaceuticals^7,17–20^. Monoclonal antibodies (mAbs), which are typically produced in mammalian cell lines due to their requirement for human-like post-translational modifications^21^, including *N*-linked glycans, can be produced in *P. tricornutum*. Strikingly, *P. tricornutum* possesses a core *N*-linked glycosylation pathway very similar to that found in humans^22–25^. One notable difference is that *P. tricornutum* decorates *N*-glycans with a core *α*(1,3)-linked fucose, similar to land plants and invertebrate species, as opposed to the *α*(1,6)-linked fucose in mammals. The *α*(1,3) linkage can be immunogenic in humans, triggering an IgE response that precludes its presence in therapeutics^26,27^. *P. tricornutum* contains three putative core fucosyltransferases (FucTs) in its genome with highly divergent nucleotide sequences.

Realizing the biotechnological potential of *P. tricornutum* requires reliable and easy-to-use gene-editing tools that enable simultaneous targeted modifications at multiple loci such as the *FucT* multigene family. Current Cas9 editing systems for *P. tricornutum* are limited by the number of single guide RNAs (sgRNAs) that can be multiplexed. Here, we develop a multiplexed Cas9 editing system for simultaneous knockout of multigene families, or of multiple genes simultaneously. Our system, PHYCUT (***<u>Ph</u>****aeodactylum tricornutum* Cs**<u>y</u>**4-**<u>C</u>**as9 m**<u>u</u>**ltiplex **<u>t</u>**ool), utilizes the Csy4 endoribonuclease to process a synthetic multi-guide array transcribed from a Cas9-expression plasmid. We find that up to 6 sgRNAs can be expressed and processed by Csy4 and use this setup to simultaneously target all three *P. tricornutum FucT* genes that perform *α*(1,3) fucosylation in the *N*-linked glycosylation pathway. Characterization of secreted proteins revealed significantly decreased fucosylation in FucT knockout strains as compared to wild type cells. Our work provides a starting point for humanizing the *P. tricornutum N*-linked glycosylation pathway and a robust, easy-to-use, multiplexed Cas9 editing system for probing *P. tricornutum* biology or for synthetic biology applications.

## RESULTS

### Processing of sgRNA arrays by Csy4

We adapted the Csy4 endoribonuclease from *Pseudomonas aeruginosa* PA14 to enable a plasmid-based multiplexed gene-editing system in *P. tricornutum* (PHYCUT) (Figure 1A, B). In the native CRISPR system, Csy4 processes pre-crRNA arrays by recognizing a 20-nucleotide sequence that forms a hairpin structure, cleaving at position G20 to liberate the downstream crRNA^28^. Csy4 can also process synthetic CRISPR arrays^29,30^. All components of the editing system were installed on the same plasmid, including codon-optimized sequences for Csy4 and *Streptococcus pyogenes* Cas9 (SpCas9). The PHYCUT vector expresses SpCas9 under the control of the inducible nitrate reductase promoter and terminator^31,32^, and Csy4 under control of the constitutive H4-1*β* promoter and terminator^5^. An sgRNA cassette was designed to contain a single sgRNA scaffold sequence adjacent to BsaI sites, enabling cloning of multiple sgRNA inserts that included the Csy4 processing motif, the spacer sequence, and sgRNA handle.

**Figure 1.**
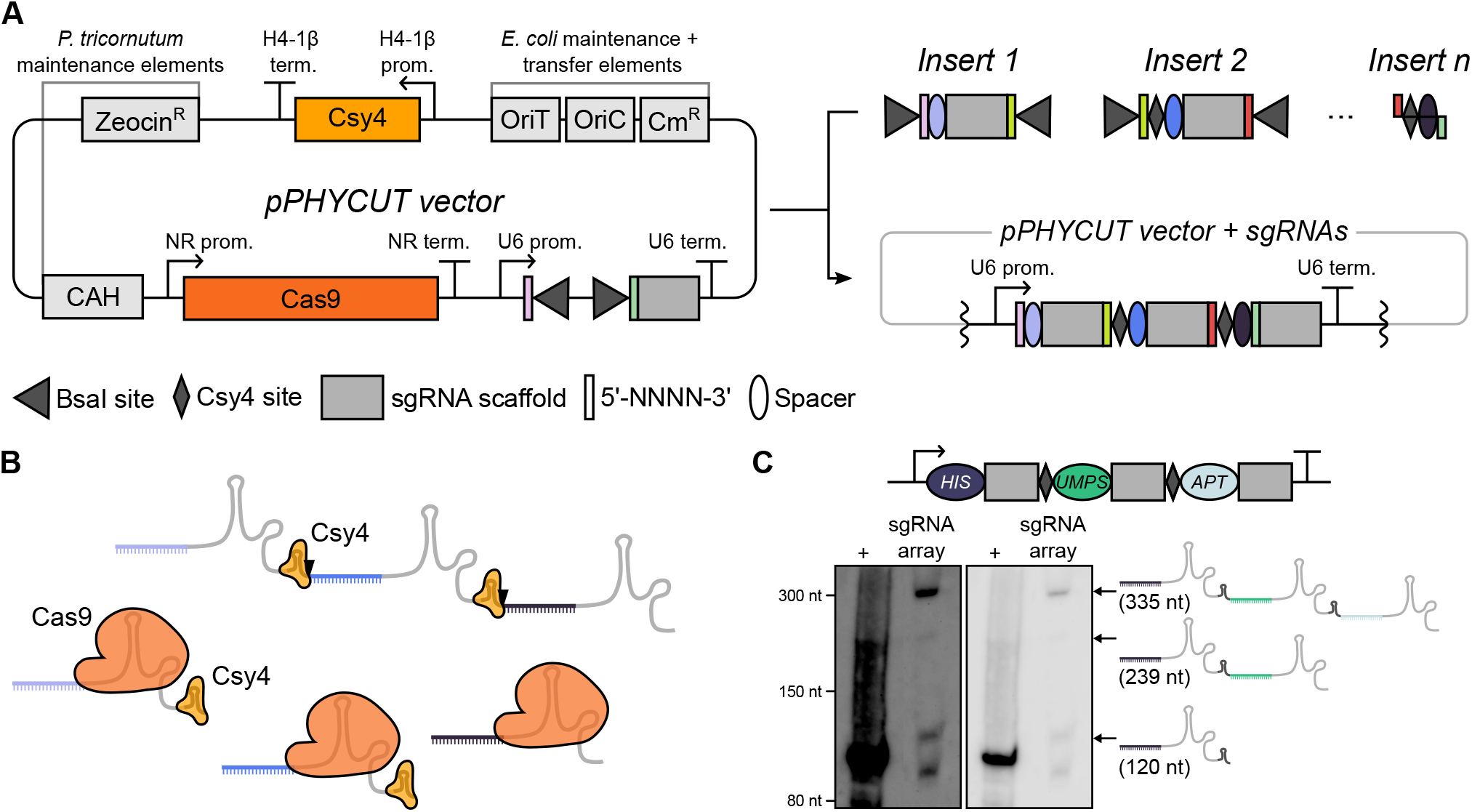
Overview of the PHYCUT editing system. A) Plasmid design and cloning strategy for the multiplexed sgRNAs. Csy4 is constitutively expressed, Cas9 is regulated by the nitrate reductase (NR) promoter and terminator, and the sgRNA array is expressed as a single transcript from the U6 promoter and terminator. Genetic elements for propagation and those required for conjugation from *E. coli* to *P. tricornutum* are shown in light grey. The sgRNA inserts include BsaI cut sites that generate unique 4nt flanking sequences to facilitate cloning. B) Schematic of Csy4-processing of a multi-guide transcript expressed from the U6 promoter. C) Northern blot of total RNA isolated from cells expressing the multi-guide transcript using a HIS crRNA-specific probe. The (+) lane is a synthesized 120-nt ssDNA analogous to the first sgRNA in the array. Identities of predicted processing products are indicated to the right. The gel image on the left shows the blot with a higher contrast.

To test the expression and processing of the multiplexed sgRNAs by Csy4, we constructed a plasmid encoding three sgRNAs previously used to target the *P. tricornutum* phosphoribosyl-ATP pyrophosphohy-drolase (*HIS*), uridine monophosphate synthase (*UMPS*), and adenine phosphoribosyltransferase (*APT* ) genes^33^ (Figure 1C). Total RNA was isolated from cells harbouring the plasmid and used for Northern blot analysis with a probe corresponding to the spacer region of the first sgRNA in the array. As shown in Figure 1C, we observed RNA species corresponding to an unprocessed array transcript as well as RNA species corresponding to processed sgRNAs, consistent with the Csy4 processing of the array. Increased contrast of the Northern blot revealed an RNA species consistent with a processing intermediate of two sgRNAs. We also noted a smaller species around 90–100 nt of unknown identity.

### PHYCUT enables multiplexed editing in *P. tricornutum*

Encouraged by the finding that Csy4 can process sgRNA arrays, we tested the multiplexing capabilities of the system by targeting the adenine phosphoribosyltransferase (*APT* ) gene, knockouts of which can be directly selected on solid agar plates supplemented with 2-fluoroadenine (2-FA)^13^ (Figure 2A). We first sought to determine whether two guides targeting the same gene would increase knock out efficiency, and designed two sgRNAs targeting the *APT* gene that were separated by ∼ 300 bps. These guides were cloned individually or as a pair into the PHYCUT construct and delivered into *P. tricornutum* by electroporation. We optimized media conditions and recovery time from electroporation for 2-FA and zeocin selection on solid media plates, finding that increased recovery times (2–3 days) relative to established electroporation protocols^9^ were necessary to recover comparable levels of electrotransformants selected on zeocin (zeo)-containing solid media versus nourseothricin selection (Supplemental Figure S1). For *APT* knock out selection, 50 *µ*M Zeocin (Zeo50) and 10 *µ*M 2-FA in plates eliminated background growth seen in electroporations, whereas 10 *µ*M 2-FA alone was sufficient to reduce background in conjugation experiments.

**Figure 2.**
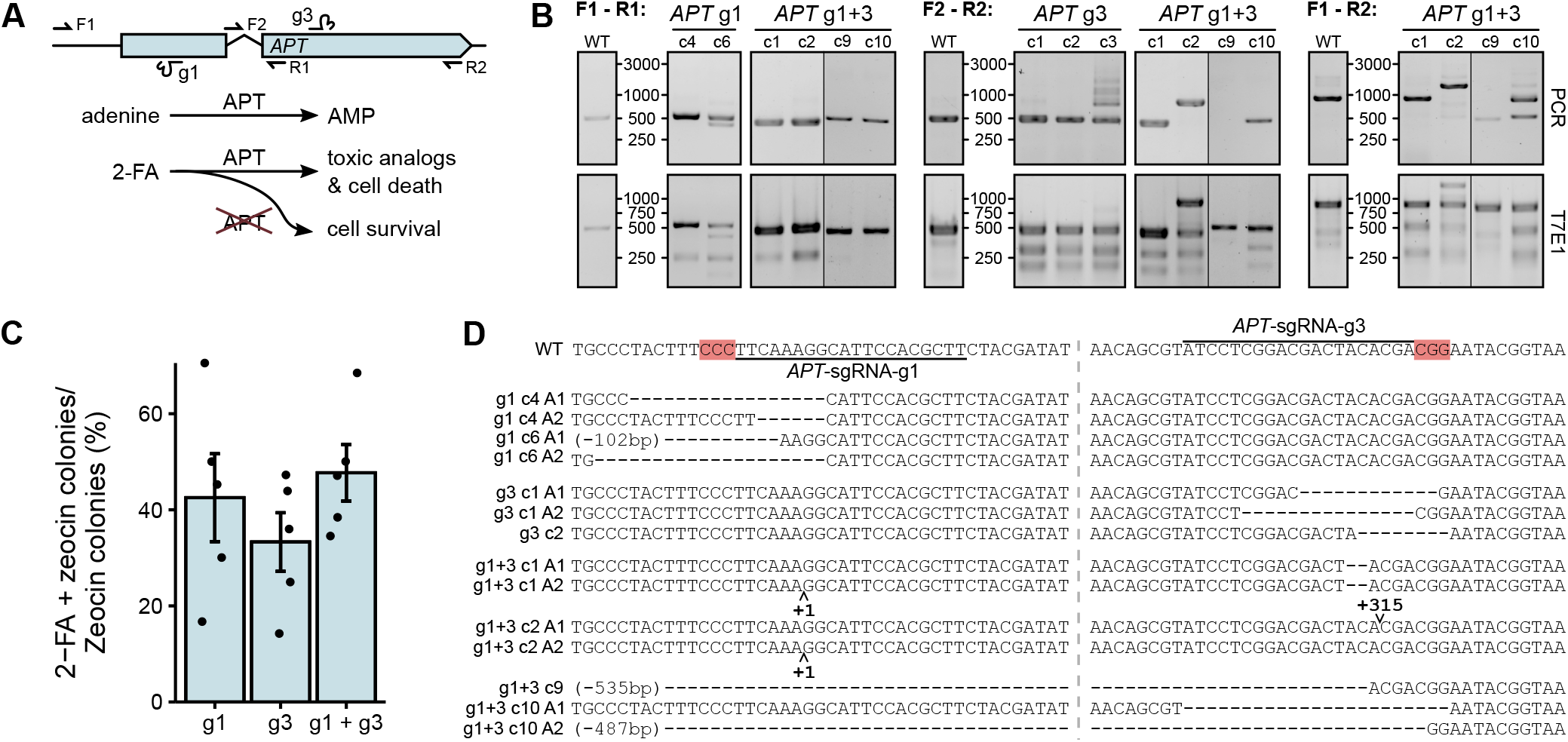
PHYCUT system testing. A) The counter-selectable *PtAPT* gene is shown with the SpCas9 target sites and screening primer locations indicated. The single intron is indicated by a gap. Consequences of an *APT* knockout in the presence of 2-FA are shown below. AMP, adenosine monophosphate. B) Representative agarose gels of PCR products (top) and T7E1 assays (bottom) to assess editing at the indicated PCR amplicons (F1-R1, F2-R2, F1-R2). Sizes (in bp) are indicated next to the gel images. WT indicates PCR amplicons and T7E1 digests from untreated *P. tricornutum*. Note that T7E1 digests contain a mixture of PCR amplicons from WT and edited strains. Vertical lines indicate cropped images from different agarose gels. C) *APT* knock out efficiency of *APT* - sgRNA-g1, *APT*-sgRNA-g3, or both, determined by the ratio of transformants on solid media containing 2-FA and zeocin, or zeocin only. Barplots are mean of 5 biological replicates with error bars representing standard error of the mean. D) Sequences of edited *APT* genes from the indicated transformants. The vertical dashed light grey line indicates the spatial separation between the two target sites. The size of deletions are indicated in parentheses for deletions extending outside of the region shown. The *APT*-sgRNA-g1 and *APT*-sgRNA-g3 sites are indicated on the WT sequence with red shading indicating the PAM motif.

The ratio of colonies on 2-FA-containing plates to colonies on Zeo-containing plates was used to measure *APT* knockout efficiency. For *APT*-sgRNA-g1 and *APT*-sgRNA-g3 expressed as single guides, we found an average *APT* knock out efficiency of 43% and 33%, respectively (Figure 2C). Expressing both guides resulted in an increased efficiency (48%) relative to the individual guides, but this increase was not statistically significant. T7 endonuclease 1 (T7E1) digests of PCR products amplified from 2-FA resistant transformants confirmed that editing occurred at the targeted loci (Figure 2B, Supplemental Figure S2A). Editing was observed at both targeted loci in 16 of 18 2-FA resistant transformants screened, indicating that both sgRNAs were active. Two exconjugants (C9 and C10, Figure 2B, *APT* g1+g3) showed evidence of large deletions that were not observed when only either sgRNA was expressed individually, suggesting that multiplexing two sgRNAs can induce large deletions between the two target sites. However, the predominant outcome were small indels at the individual sites.

Additionally, we designed an sgRNA (*APT*-sgRNA-g2) that targets a site containing a single nucleotide polymorphism that results in either a NGG or NGA PAM, based on previous reports that SpCas9 can target NGA PAM sites^34^. This guide, as well as *APT*-sgRNA-g1, were cloned individually or together and delivered into *P. tricornutum* via bacterial conjugation from *E. coli* (Supplemental Figure S2B,C). We found a much lower editing rate for *APT*-sgRNA-g2 as determined by 2-FA resistant exconjugants (5%) and T7E1 digests of PCR amplified samples. Sanger sequencing showed that *APT*-sgRNA-g2 target sites with NGA PAM sequences were not substrates for SpCas9 editing, whereas sites with NGG PAMs showed evidence of editing (Supplemental Figure S2D). When multiplexed with *APT*-sgRNA-g1, we found an increase in knock out efficiency (19%) compared to either individual sgRNA, but this increase was not found to be significant compared to *APT*-sgRNA-g1 (16%). Notably, the knockout efficiency as determined by selective colony count ratios was over 2-fold greater (43% vs. 16%) for *APT*-sgRNA-g1 when electroporated into *P. tricornutum*, versus delivery by conjugation (Figure 2C, Supplemental Figure

S2C). This difference could reflect past observations that ∼ 10–20% of *P. tricornutum* exconjugants have rearrangements in conjugative plasmids^8,35^, which in this case could impact PHYCUT function.

### Guide order within the array influences editing efficiency

We next tested whether the linear order of sgRNAs in the array impacted editing efficiency of sgRNAs, possibly because processing of the primary guide RNA transcript by Csy4 would be incomplete. We designed an array consisting of *APT*-sgRNA-g1 and a previously used sgRNA targeting the *UMPS* gene (*UMPS*-sgRNA)^33^ in either position of the array (*APT* |*UMPS* or *UMPS* | *APT* ) (Figure 3A, B). Knockouts of the *UMPS* gene are uracil auxotrophs and are resistant to the nucleotide analog 5-fluoroorotic acid (5-FOA)^33,36^. Knock out efficiency of *APT* was determined by 2-FA resistant exconjugants for both constructs (Figure 3C). Subsequently, 2-FA resistant exconjugants were grown on plates containing

**Figure 3.**
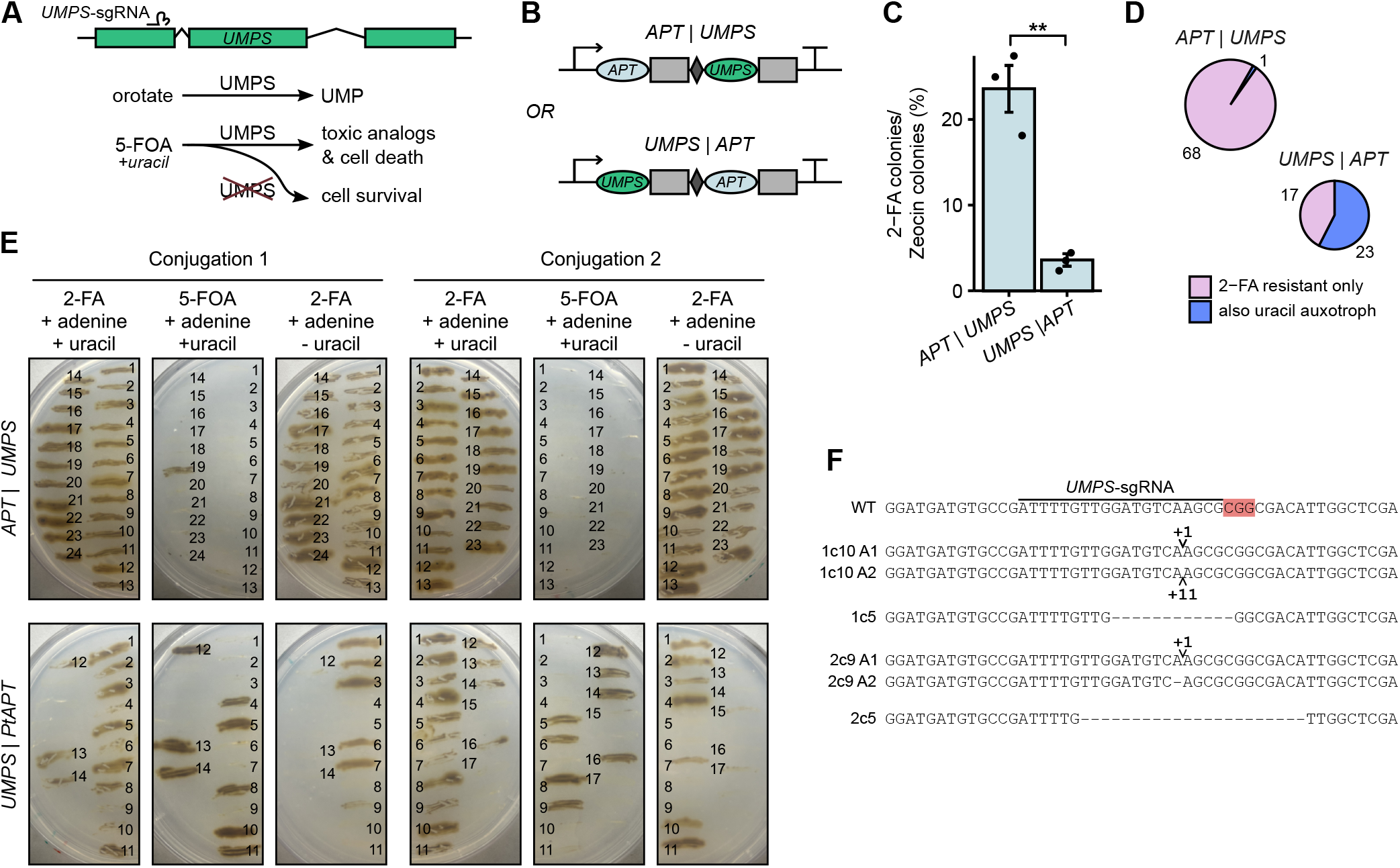
PHYCUT guide order testing. A) The counter-selectable *PtUMPS* gene is shown with the SpCas9 target site indicated. The two introns are indicated by gaps. Consequences of an *UMPS* knockout in the presence of 5-FOA are shown below. UMP, uridine monophosphate. B) The two guide arrays used to determine linear order effects, expressing *APT*-sgRNA-g1 and *UMPS*-sgRNA. C) *APT* knock out efficiency of the two arrays in panel (B), determined by the ratio of exconjugants on solid media containing 2-FA or zeocin. Barplots are a mean of 3 biological replicates with error bars representing standard error of the mean. **, p *<* 0.01. D) *UMPS* knock out efficiency for dual-gene array exconjugants selected on 2-FA, as determined by re-streaking exconjugants onto plates containing 5-FOA and uracil. E) Streaks of the dual-gene array exconjugants selected on 2-FA, growing differentially on plates with or without 5-FOA or uracil supplementation. F) Sequences of the edited *UMPS* gene from indicated exconjugants. A 1 or 2 indicates the conjugation experiment, followed by the clone (c) number, and the different alleles (A1 or A2) if the edits were not homozygous. The *UMPS*-sgRNA site is indicated on the WT sequence with red shading indicating the PAM motif.

FOA, or plates lacking uracil supplementation, to determine whether they were also *UMPS* knock outs (Figure 3D,E). We found that when *APT*-sgRNA-g1 was first in the array (*APT*| *UMPS*), 24% of exconjugants were 2-FA resistant, but only 1 of 69 tested exconjugants was also resistant to 5-FOA (Figure 3C,D). Conversely, when the *UMPS*-sgRNA was first in the array (*UMPS*| *APT* ), a lower rate of 2-FA resistant exconjugants were found (4%), but more than half of exconjugants screened were both *UMPS* and *APT* knockouts (23 of 40 tested). Sanger sequencing of edited *UMPS* genes from exconjugants receiving the *UMPS* | *APT* construct revealed both insertions and deletions localized to the sgRNA target site (Figure 3F). This data suggests that position in the array rather than intrinsic sgRNA activity accounts for the observed differences in editing efficiency.

### PHYCUT can multiplex six sgRNAs for editing

To determine if the PHYCUT array could express more than two sgRNAs, we designed arrays that contained the *APT*-sgRNA-g1 and g3 guides in the distal positions of the array relative to the promoter that were tiled out with one, two, three or four non-targeting guides in the proximal positions (Figure 4A). The non-targeting guides had at least 5 mismatches to the *P. tricornutum* genome sequence. These constructs were then electroporated into *P. tricornutum*, and editing was assessed as before. We found that increasing the array size from two sgRNAs to four sgRNAs did not significantly decrease *APT* knock out efficiency (48% for two, 46% for three, 41% for four). At five and six sgRNAs, efficiency was reduced to 7% and 9%, respectively. However, even at a length of six sgRNAs editing is clearly observed (Figure 4B). Transformants 2 and 3 (indicated by # and †, respectively) that received the six-sgRNA construct showed the presence of an amplicon that indicates a large deletion between the two *APT* guide target sites, deleting the F2 and R1 primer binding sites. This deletion was found to be homozygous in transformant 3, where no amplicon could be seen with the F1-R1 or F2-R2 primer pairs, and heterozygous for transformant 2, where the other allele possessed an indel at the *APT*-sgRNA-g1 site. However, for four of the eight six-sgRNA transformants screened, editing was only detected at the *APT*-sgRNA-g1 locus and not at the g3 site (Figure 4B), in contrast to the comparable levels of editing seen for each guide in a two-sgRNA array (Supplemental Figure S2A). This indicates that as sgRNA array size increases, activities of distal sgRNAs beyond a length of 4 decreases.

**Figure 4.**
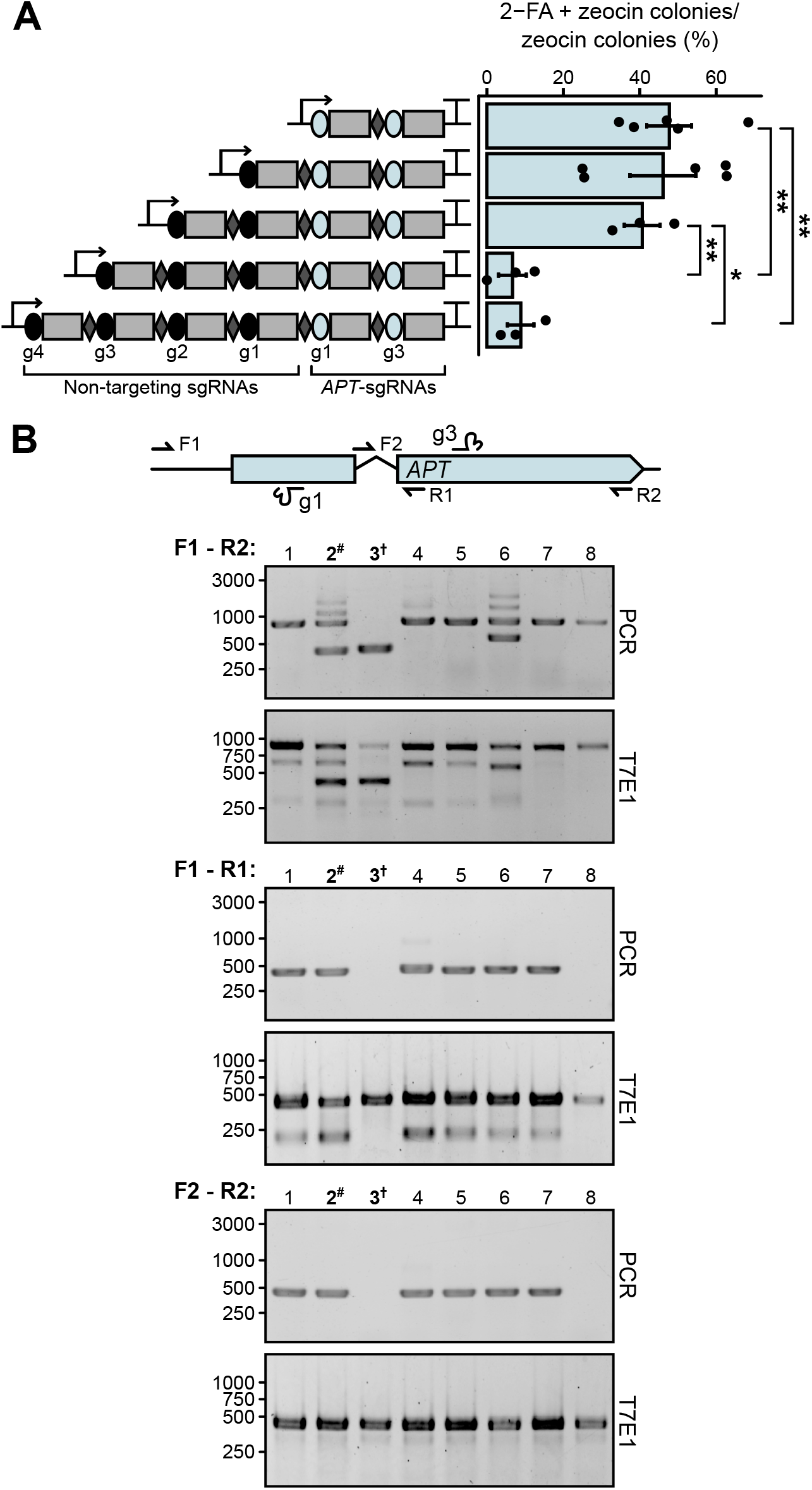
Testing sgRNA array length. A) On left, Schematic of the tiled *APT*-sgRNA-g1 and g3 constructs, labeled as in Figure 1A. Note that *APT*-sgRNA-g1 and g3 are in the distal positions of the array relative to the promoter. On right, knockout efficiency of each array as assessed by 2-FA resistant transformants. Barplots are a mean of 3–5 biological replicates with error bars representing standard error of the mean. Significant changes in knockout efficiency between the different constructs are displayed, with * p *<* 0.05, ** p *<* 0.01. Comparisons not shown are non-significant. B) Screening of editing events in transformants that received the six-sgRNA construct. Locations of screening primers are indicated. For each primer combination, an agarose gel image of the PCR prdocuts and T7E1 digests are shown for the indicated transformants. Sizes (in bp) are indicated next to the gel images. T7E1 digests contain a mixture of PCR amplicons from WT and edited strains. Note that transformant 3 (†) has a large homozygous deletion and cannot be amplified by primer sets F1-R1 or F2-R2. Transformant 2 (#) is heterozygous and shows editing by *APT*-sgRNA-g1 and g3 in one allele, and by g1 only in the other allele.

Collectively, this data demonstrates that a plasmid-encoded multiplexed sgRNA array can be processed by Csy4 to enable SpCas9 editing at multiple sites in the *P. tricornutum* genome. Our testing of two sgRNAs targeting different genes shows that the order of sgRNA in the array impacts editing efficiency, although intrinsic sgRNA activity could also influence editing efficiency. We also showed that the PHYCUT system can express six sgRNAs. Editing efficiency is reduced for arrays with sgRNAs in the fifth and sixth positions. Further, our data indicates that electroporation of PHYCUT constructs rather than delivery by conjugation can increase editing efficiency.

### RNAseq of the *N*-linked glycosylation pathway

The predicted *N*-linked glycosylation pathway of *P. tricornutum* is similar to that of humans^22,23^ and multiplexed Cas9 editing tools would aid in enabling the full humanization of this pathway. As a prelude to further glycoengineering of *P. tricornutum*, we used RNAseq to examine the expression of genes with predicted functions in the *N*-linked glycosylation pathway (Figure 5A,B and Supplemental table S1). We were particularly interested in the *P. tricornutum* FucT (PtFucT) enzymes, as these represent a target for multiplexed CRISPR knockouts to remove the *α*(1,3)-linked fucose from recombinant proteins purified from *P. tricornutum* that would otherwise be immunogenic in humans^26,27^. *P. tricornutum* encodes three CAZy GT10-family FucTs (PtFucT1, PtFucT2, and PtFucT3). These FucTs contain the well-conserved C-terminal GT10 domain^37^ responsible for donor GDP-fucose binding^38^, but the N-terminal domains responsible for acceptor-substrate binding are not as conserved, as is the case with many plant FucTs.

**Figure 5.**
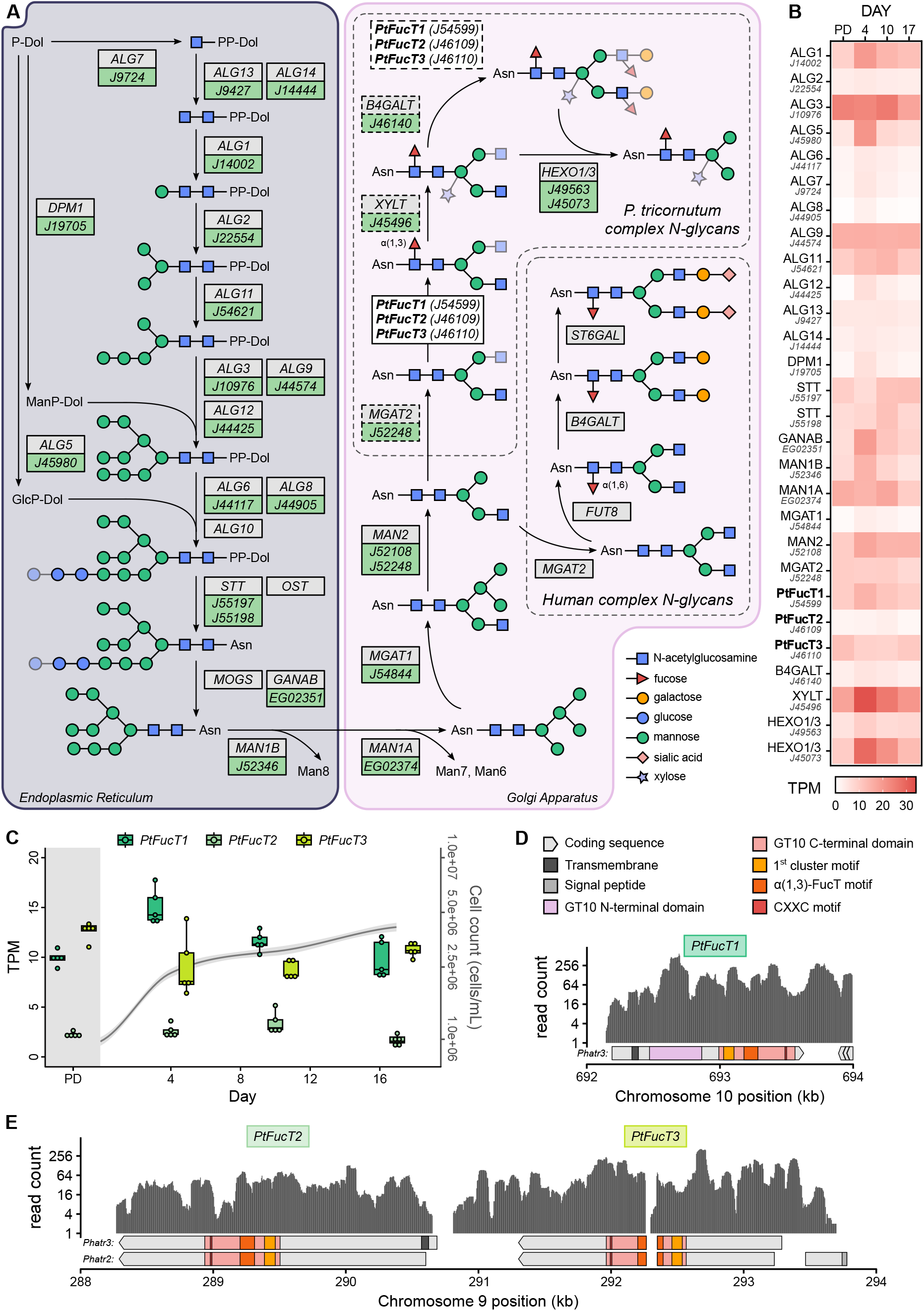
RNA-seq analysis of the *P. tricornutum N*-linked glycosylation pathway. A) Predicted *N*-linked glycosylation pathway of *P. tricornutum*. Grey boxes contain the KEGG name for glycosyltransferase enzymes, light green boxes contain the Phatr3 annotation names for *P. tricornutum* orthologs. Dashed boxes and translucent moieties show reactions that lack experimental evidence supporting their presence or precise location in *P. tricornutum*. The human complex N-glycan shows a sialylated biantennary structure as an example, which are the predominant glycoform for mAbs. B) RNA-seq reads converted to transcripts per kilobase million (TPM) for glycosyltransferase enzymes in wildtype *P. tricornutum* across four different timepoints. PD = pre dilution, sample taken immediately before diluting cultures for time course. C) TPM expression levels of the three PtFucTs across four different time points in wildtype *P. tricornutum*. A loess smooth regression line for the cell densities of the five replicate cultures is plotted on the secondary y-axis on a log10 scale in grey, with the 95% confidence interval shown in shaded light grey. D) Read coverage for the three PtFucTs, plotted on a Log2 scale and separated into 5nt bins. The PtFucT coding sequences annotated for both Phatr2 and Phatr3 are shown below, with predicted FucT domains and motifs included.

Acceptor-substrate specificity of the three PtFucTs remains ambiguous, with them potentially adding the *α*(1,3)-fucose moiety to the chitobiose core^22,37^, a terminal GlcNAc branch as in Lewis-epitope formation^39^, or even to the core mannose residues^40^. Regardless, these inappropriate additions must be attenuated to achieve human-like complex *N*-glycans.

Phylogenetic analyses revealed that PtFucT2 and PtFucT3 cluster in a divergent clade of SAR-specific FucTs, with the diatom FucT2 potentially resulting from a gene duplication event of FucT3 that occurred after the divergence of pennate diatoms (Supplemental Figure S3, Supplemental Table S2). Interestingly, PtFucT1 is nested in a clade of arthropod-specific FucTs with very few pennate diatom-specific orthologs, indicating a more recent horizontal gene transfer (HGT) event in a pennate diatom ancestor. This finding is consistent with phylogenetic analyses showing that the *P. tricornutum* genome is highly enriched in HGT events from bacterial and green algal species^3,41,42^. The sequence of PtFucT1 predicts a protein organization very similar to invertebrate-type *α*(1,3)-FucTs, a group which comprises both core and Lewis-type FucTs.

Examining the expression levels of the three *PtFucT* genes more closely, we noted that *PtFucT1* and *PtFucT3* show comparatively higher levels of expression throughout the growth cycle compared to *PtFucT2*, which shows lower expression levels in the range of ∼ 1.5–3.5 TPM (Figure 5C). We examined the RNAseq read coverage for the three *PtFucTs* to determine the exonic boundaries of the genes and to help resolve discrepancies in the predicted 5’ ends of *PtFucT2* and *PtFucT3* from different genome annotations (Figure 5D, E, Supplemental Table S3)^3,41^. We found that reads mapped to the 5’ end of *PtFucT2* corresponding to the Phatr2 annotation^3^ rather than extending upstream, indicating that for our strain and culturing conditions, the Phatr2 annotation for *PtFucT2* may be more accurate. For *PtFucT3*, the mapped reads were consistent with splicing of the intron between exon 2 and 3 (Phatr2 annotation), although a small proportion of reads aligned within the 5’ end of the intron. This sequence contains two stop codons, suggesting there may be a regulatory aspect to retention of this intron. We did not find reads mapping to the upstream region of exon 1 based on the Phatr2 annotation. The mapped reads for *PtFucT3* also support the possibility of an alternative start codon, 222 nt upstream and in-frame of the Phatr3 annotation start. This PtFucT3 707 aa isoform includes a transmembrane domain that is predicted to localize to the Golgi apparatus. The *PtFucT1* annotation remains unchanged between Phatr2 to Phatr3 and also encodes a transmembrane domain for Golgi localization and a protein architecture similar to invertebrate FucTs.

### PHYCUT-generated knockouts of the FucT multi-gene family

The coding DNA sequences of the three FucT paralogs are sufficiently divergent such that a single, common sgRNA cannot target all three genes. A PHYCUT construct was designed to express three different sgRNAs with each sgRNA targeting one of the FucT paralogs (Figure 6A,B, Supplemental Table S4). Of note, the sgRNAs were designed based off the Phatr2 coding sequence annotations, thus the *PtFucT3* guide targets a region upstream of the translational start site predicted by the Phatr3 annotation. After conjugation of the PHYCUT plasmid from *E. coli* to *P. tricornutum*, exconjugants were selected on zeocin-containing plates and screened for editing using T7E1 assays of PCR products corresponding to each targeted loci (Figure 6C, Figure S4A). Long-range PCRs were also conducted to screen for larger deletions occurring between the two adjacent sgRNA target sites in *PtFucT2* and *3* (Figure 6D, Figure S4B,C). Of the 30 primary exconjugants screened, 15 showed editing at all three targeted loci, with only one exconjugant having no editing detected at any loci (Figure 6E). We also observed defined-length deletions between the *PtFucT2* and *3* target sites in 8 of the 30 exconjugants screened, as well as other larger deletions occurring at all three targeted locations. Because exconjugant colonies contain a population of different editing events, we chose four exconjugants (2, 5, 21, and 27) that clearly showed editing at the three *PtFucT* genes and isolated 15 subclones from each of them. We found that between 7-53% of subclones showed editing of the three *PtFucT* genes, depending on the subclone population (Figure 6E, Supplemental Figures S5 and S6). At the same time, we used a non-multiplexed CRISPR-Cas9 plasmid to target the *PtFucT2* and *PtFucT3* genes individually (Supplemental Table S4). Sanger sequencing was performed to determine genetic changes from editing (Figure 6F, Supplemental Tables S5 and S6); notably, we were unable to identify any strains harbouring a biallelic deletion of all three *PtFucT* copies. We selected 25 strains for further phenotypic characterization, 18 from the PHYCUT multiplex knockouts and 7 from single sgRNA knockouts.

**Figure 6.**
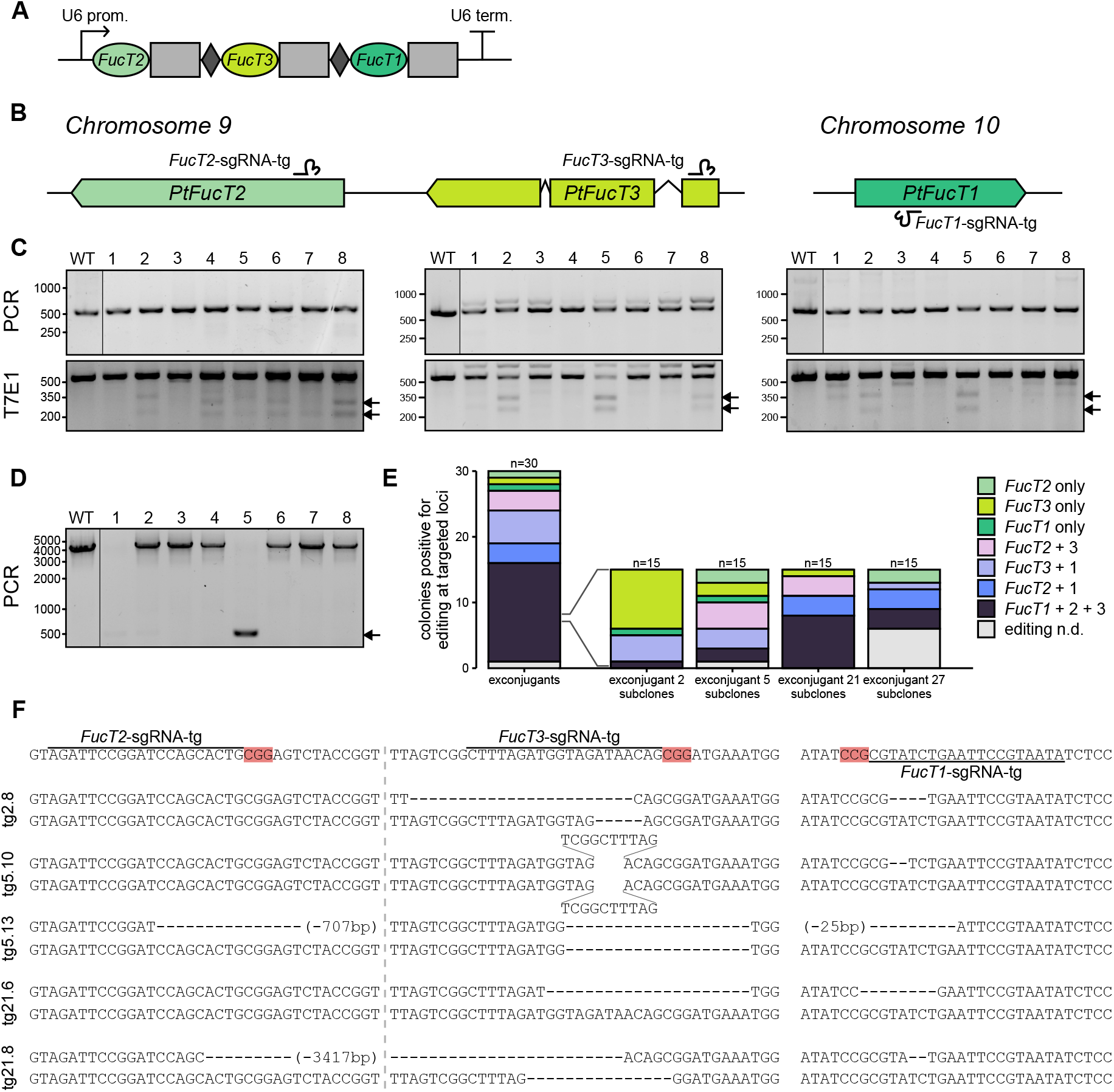
PHYCUT knockout of three *FucT* genes in *P. tricornutum*. A) PHYCUT sgRNA array expressing the three *FucT* guides. B) The three *P. tricornutum FucT* genes according to the Phatr2 annotations with their guide target locations shown. C) Short-range PCRs and T7 Endonuclease (T7E1) assays of the three target genes for representative exconjugants. D) Long-range PCR surrounding target sites for *PtFucT2* and *3* show large deletions between the target sites. The amplicon size of a deletion from the sgRNA-*PtFucTA*-tg to sgRNA-*PtFucTB*-tg target sites is 414bp. E) Proportion of colonies screened that showed editing events at each of the three loci, as determined by PCRs and T7 assays. Four exconjugants showing evidence of editing at all three loci were replated for subclones and re-screened. F) Sanger sequencing data of selected subclone strains at each of the targeted FucT loci. The dashed light grey line indicates the spatial break between the PtFucTA and B target loci. For deletions spanning regions larger than the sequence shown, the total deletion size is indicated in parentheses.

### FucT knockouts show decreased core *α*(1,3)-fucosylation of secreted proteins

To assess whether the *PtFucT* knockouts would lower the levels of core *α*(1,3)-fucosylation, we developed an indirect ELISA protocol that utilized an antibody specific to the core *α*(1,3)-fucose epitope to detect this moiety in secreted proteins (Supplemental Figure S7A-C). We hypothesized that secreted proteins that have fully transited throughout the secretory system should contain the highest proportion of matured, and therefore fucosylated glycoproteins, allowing us to detect fucosylation changes in the mutant strains as compared to wild type. The 25 strains selected for further analysis, along with three wild-type *P. tricornutum* cultures grown independently for at least 2 months prior to the experiment, were grown over a course of 12 days and changes to fucosylation profiles assessed on days 4, 8, and 12 (Figure 7A, Supplemental Figures S7-9). For wildtype *P. tricornutum*, we found that the three cultures exhibited varying degrees of fucosylation, with between 1 x 10^9^ - 8 x 10^9^ fucose sites per ng of protein. This indicates natural variation in fucosylation levels of glycoproteins between different clonal populations of *P. tricornutum* (Supplemental Figure S7D, E). We noted that total fucosylation increased in secreted proteins from day 4 to day 8 and remained high on day 12. Normalizing fucosylation levels to extracellular protein concentration revealed a peak on day 8 and a decrease to day 12. This may indicate that in later growth stages cells are secreting a lower proportion of fucosylated glycoproteins, possibly due to the lower expression of MGAT1 measured by RNAseq in later growth stages (Figure 5B).

**Figure 7.**
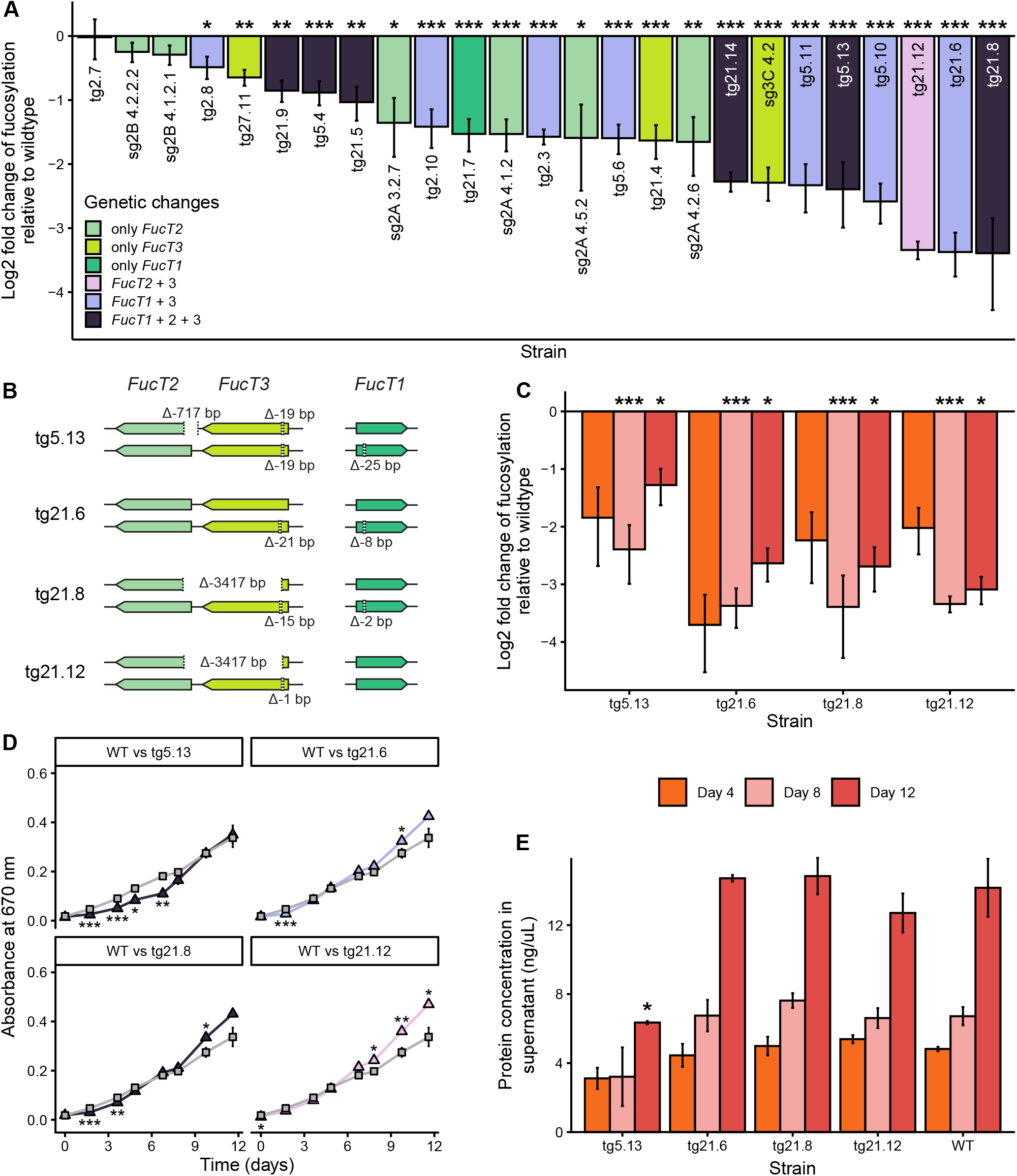
FucT mutants showed decreased core-fucosylation of secreted proteins. A) *α*(1,3)-fucosylation levels of secreted proteins for FucT knock out strains on day 8 normalized to their culture density and plotted as log2 fold change relative to wild type. Strains are coloured by their genetic edits. P-values compared to the wild type average are displayed above each sample. * p *<* 0.05, ** p *<* 0.01, *** p *<* 0.001. B) Genetic changes of four mutant strains that showed persistently low fucosylation across all three time points. C) *α*(1,3)-fucosylation of glycoproteins secreted into the extracellular media by the four persistently low FucT mutant strains normalized to their culture density, plotted as log2 fold change relative to wild type *P. tricornutum*. P-values compared to the wild type average for each respective day are displayed above each sample. D) Growth curves of the four persistently low FucT mutant strains (triangles) compared to wildtype *P. tricornutum* (grey squares), as determined by absorbance at 670 nm. Strains are coloured by their genetic edits as in panel (A). E) Concentration of proteins in the extracellular media of the four persistently low FucT mutant strains and wildtype *P. tricornutum*. P-values compared to the wildtype average for each respective day are displayed above each sample. All plots show the average of three biological replicates, with error bars representing standard error from the mean.

Decreases in fucosylation compared to the wildtype averages were seen for the FucT mutant strains across all three time points measured, with the average log2 fold change being -1.66, -1.61, and -1.16, for days 4, 8, and 12, respectively (Figure 7A and Figure S7-9). The largest log2 fold change seen for the three time points for any of the samples were -3.70, -3.39, and -3.09, by the strains tg21.6 on day 4, tg21.8 on day 8, and tg21.12 on day 12. The PHYCUT-edited strains showed larger decreases in fucosylation on average than strains generated by a single-guide construct, most notably on day 4 where PHYCUT-edited strains showed a 75% decrease in fucosylation as compared to the single-guide cohort (p *<* 0.0001) (Figure 7A). Among the strains evaluated, four (tg5.13, tg21.6, tg21.8, tg21.12) consistently showed a greater than average decrease in fucosylation across all three time points measured (Figure 7C). These strains were all edited with the multiplexed PHYCUT construct and contained edits in at least two of the three FucT genes (Figure 7B). The largest difference in fucosylation was seen on day 8, with these four strains showing an average decrease of 88% relative to wild-type cells (p *<* 5x10^-6^). These four strains grew comparably to the wildtype controls, with tg21.6, tg21.8, and tg21.12 potentially growing better than the wild type in later stages (Figure 7D). A small growth defect was seen for tg5.13 in earlier time points, however this difference was eliminated from day 8 onwards Figure 7D, Supplementary Figure S10). Levels of secreted protein were not impacted for strains tg21.6, tg21.8, and tg21.12, but strain tg5.13 showed a marked decrease in extracellular protein levels on day 12 (Figure 7E). Collectively, this data demonstrates that FucT mutant strains generated with the PHYCUT construct show significantly decreased levels of fucosylated glycoproteins, without having a negative impact on strain fitness.

## DISCUSSION

The global market for protein biologics, diagnostics and value-added chemicals is rapidly growing^43–46^.

*P. tricornutum* has the flexibility to make *N*-glycosylated biologics appropriate for human therapies such as monoclonal antibodies in a carbon-negative manner and is an excellent candidate for a scalable photosynthetic protein production platform orthogonal to mammalian or other systems. To harness the full potential of *P. tricornutum* for these biotechnological applications, reliable methods for multiplexed genome engineering are required.

Current strategies for multiplexing CRISPR/Cas9 sgRNAs in *P. tricornutum* include plasmid-based and DNA-free systems. For plasmid-based editing, two to three cloned sgRNAs transcribed by multiple independent U6 promoter/terminator expression cassettes in tandem permit transcription of sgRNAs with a defined 5’ end^47,48^. Increasing the number of sgRNAs in these cassettes may prove challenging, as direct sequence repeats can pose issues for cloning and stability of constructs. Taparia and co-workers^49^ developed a multiplexed editing system transcribed by RNA polymerase II and processed by endogeneous ribonucleases, although the accuracy of processing and guide functionality was confounded by targeting of a single gene by the multiplexed guides. DNA-free biolistic delivery of Cas9/sgRNA RNPs relies on the simultaneous knock out of counter-selectable genes such as *APT* or *UMPS*^13^, creating auxotrophic strains that may be undesirable for downstream purposes, and that limit the ability to perform successive rounds of knockouts. To address these shortcomings, we developed the plasmid-based PHYCUT system, a method of multiplexing that is easily clonable, capable of expressing up to six guides that can efficiently knock out multiple genes, and is compatible with more accessible modes of DNA delivery like bacterial conjugation and electroporation. We envision that PHYCUT plasmids could be adapted for transcriptional knockdown applications with dCas9, or other CRISPR-based editors, and cured from edited strains by simple passaging in antibiotic-free media^33^.

Northern blot analyses confirmed that Csy4 was able to process full length multiplexed transcripts into individual sgRNAs. Notably, we found that the transcript was detected as a fully processed or unprocessed species, with minimal detection of an intermediate 2-sgRNA species. This observation is consistent with biochemical characterization of Csy4 as a single-turnover enzyme^50^, suggesting that in the PHYCUT setup stoichiometric amounts of Csy4 must be present for full transcript processing.

We did not find a significant improvement in knock out efficiency when two sgRNAs were used to target the *APT* gene compared to either one individually. While editing was detected at both loci in the majority of cases, small indels at each target site were the predominant editing outcome. Interestingly, we noted a significantly higher proportion of defined-length deletions between the *PtFucT2* and *3* target sites, suggesting that large deletion efficiency could be modulated by adjusting the relative spacing or the strand targeted by each sgRNA. Expression of a construct targeting both the *APT* and *UMPS* genes confirmed that multiple genes could be knocked out at once, but also suggested a strong positional effect for order of sgRNAs in the array, with the downstream guide found to be significantly less active than the first position. However, this effect was not seen in the *APT*-sgRNA-g1 and g3 construct, where editing was seen at both target loci at roughly the same frequency. This could be due to differences in delivery method, where the *APT* and *UMPS* targeting constructs were delivered via conjugation, and the *APT*-sgRNA-g1 and g3 and guide tiling constructs were delivered via electroporation. Indeed, it has been reported that rearrangements and deletions are introduced during the conjugation process from *E. coli* to *P. tricornutum*^8,35^, which may impair PHYCUT function. Even with expression of a single sgRNA, we found higher editing rates for electroporated plasmids, with *APT*-sgRNA-g1 showing a greater than 2-fold increase in knock out efficiency compared to the same construct delivered via conjugation. We therefore suggest electroporation as the optimal method of delivery for multiplexed editing. Using electroporation, we were able to detect high levels of editing for arrays up to four sgRNAs in length, with a drop in efficiency when five or six sgRNAs were present. This could be due to size constraints of the transcript length imparted by RNA pol III, which normally transcribes short non-coding RNAs less than ∼ 500 nt^51^. In the future, combating this size limit could be tested by replacing the U6 promoter and terminator with RNA polymerase II transcriptional regulation and including an additional Csy4 recognition site at the proximal end of the array to ensure a defined 5’ end of the first sgRNA.

Using the PHYCUT construct expressing three sgRNAs targeting the three *FucTs* in *P. tricornutum*, we were able to create a selection of different FucT mutants showing significantly decreased levels of core *α*(1,3)-fucosylation of secreted glycoproteins compared to the wild type with no noticeable growth defects. Notably, we were not able to identify a strain with biallelic deletions in all three copies, which may indicate a selective disadvantage to biallelic knockouts. Indeed, a complete knock out of *PtFucT1* was recently found to show a significant growth defect compared to wild type cells^39^. However, complete abolition of FucT and XylT action has been found to be possible in the plant species *Arabidopsis thaliana*^52^ and *Nicotiana benthamiana*^53^. It is possible that delivery of the PHYCUT construct by electroporation instead of conjugation to improve multiplexed editing efficiency could generate complete knock outs.

The development of our high-throughput screening method utilizing indirect ELISAs for the detection of fucosylated glycans greatly aided our ability to characterize knock out strains. This method could be re-tooled by swapping out the anti-*α*(1,3)-fucose antibody for other antibodies or lectins for other glycoengineering applications. Ultimately, we found that even though not all copies were deleted, we were able to identify strains showing up to a 90% decrease in *α*(1,3)-fucosylation of secreted glycoproteins compared to the wild-type strain. This represents an excellent starting point for further humanization of *P. tricornutum*’s *N*-linked glycosylation pathway. Phylogenetic analyses of the *P. tricornutum* FucTs provided insight into the diversity and origins of these three genes, with RNAseq analysis allowing further characterization of their expression levels and transcriptional boundaries. In the future, the PHYCUT tool could be used to further elucidate the roles of each individual *PtFucT* by combinatorially knocking out the different copies.

In conclusion, we have designed a robust system for multiplexed editing of *P. tricornutum* with PHYCUT that is easily customized and delivered, and that can multiplex more guides than is currently possible with other editing approaches. We envision several wide-ranging applications for the PHYCUT tool with active Cas9 enzymes for knockouts or inactive dCas9 variants for transcriptional modulation. Further glycoengineering of the *N*-glycosylation pathway could be achieved by knocking out the XylT gene, which adds xylose epitopes to *N*-glycans that are also foreign to humans, and the HEXO1/3 genes, which could prevent degradation of the terminal GlcNAc residues, allowing complex biantennary structures to be built. Aside from humanization of the *N*-glycan structures, other PHYCUT applications for biologic-producing strains could include knocking out proteases. *P. tricornutum* encodes a large number of proteases, often secreted into the extracellular media^54,55^. Targeting these proteases with PHYCUT could greatly improve yields of proteins purified from cell-free media, like monoclonal antibodies^56^ or viral proteins for diagnostics^7^. The study of multi-protein families, such as the large number of reverse transcriptases encoded by *P. tricornutum*^3,54^, could be greatly helped by combinatorial knock outs made with PHYCUT.

## Materials and Methods

### Strains and growth conditions used

*E. coli* (Epi300, Epicentre) was grown in Luria Broth (LB) supplemented with appropriate antibiotics (chloramphenicol (25 *µ*g/mL) or gentamicin (20 *µ*g/mL )). *P. tricornutum* (Culture Collection of Algae and Protozoa CCAP 1055/1) was grown in modified full aquil salt or half aquil salt L1 medium, as previously described^7^ at 18^*°*^C under cool white fluorescent lights (75 *µ*E) and a photoperiod of 16 h light:8 h dark. Solid media plates were made by mixing equal parts L1 or half salt L1 with 2% agar (BD BACTO™ Agar) and adding appropriate antibiotic selection. Plasmid uptake was selected for on 25% salt L1 solid media with 100 *µ*g/mL zeocin for conjugation experiments, and 50% salt L1 solid media with 50 *µ*g/mL zeocin for electroporation experiments. *APT* knock out selection was performed on 50% salt L1 solid media with 5 mg/L adenine and 10 *µ*M of 2-fluoroadenine (2-FA) as previously described^13^ for conjugation experiments, and additionally supplemented with 50 *µ*g/mL zeocin for electroporation experiments. *APT* knock out selection plates for experiments with an *UMPS*-targeting sgRNA were also supplemented with 50 *µ*g/mL uracil. To test strains for *UMPS* knock outs, streaked out exconjugants on *APT* knock out selection plates were re-patched onto 50% salt L1 solid media containing 150 *µ*g/mL 5-fluoroorotic acid (5-FOA), 50 *µ*g/mL uracil, and 5 mg/L adenine, or 2-FA and adenine plates without uracil supplementation.

### Plasmid design and construction

A list of plasmids, DNA constructs, and DNA oligonucleotides used in this study and those used to make the pPHYCUT and pSS29 plasmids are described in supplemental tables S7, S8, and S9, respectively. Briefly, pSS29 was create by swapping out the origin and *E. coli* antibiotic selection marker of pPtGE34 with pACYC-duet, and swapping the *FcpB* promoter from *SpCas9* with the *Nitrate reductase* (*NR*) promoter. pPHYCUT was then built from pSS29, cloning in the *P. tricornutum* codon-optimized *Csy4* from *P. aeruginosa* under control of the *H4-1* promoter and terminator, which was domesticated to remove BsaI sites. All plasmids were cloned with the NEBuilder® HiFi DNA Assembly Master Mix (New England Biolabs) with equimolar amounts of pieces, according to manufacturer’s instructions. PCR amplification of pieces was performed with PrimeSTAR GXL DNA Polymerase (Takara) using the standard or rapid manufacturer’s protocols. After assembly, plasmids were sequence verified using full plasmid sequencing from Flow Genomics. Cloning of sgRNAs into the Cas9 constructs was performed as described^57^ using the 3+ piece assembly protocol, and transformed into *E. coli* for screening. sgRNA inserts were purchased as g-blocks from IDT, generated via second-strand synthesis reaction from overlapping oligos, or phosphorylated and annealed from overlapping oligos in the case of the last sgRNA of the arrays. Second-strand synthesis of oligos was carried out by combining 0.75 *µ*M each of top and bottom strand oligos with 2 *µ*L NEB2 buffer and 14.2 *µ*L ddH_2_O, then incubating at 95^*°*^C for 5 minutes and slowly cooling to 4^*°*^C. 0.4 *µ*L of 2.5 mM dNTPs and 0.4 *µ*L of Klenow fragment (New England Biolabs) were added and the reaction was incubated for 30 minutes at 37^*°*^C. Reactions were then cleaned up over a spin column (HiPure DNA Kit, GeneBio Systems). *E. coli* transformants were colony screened for array insertion, and cloned plasmids were Sanger sequenced at the London Regional Genomics Centre to verify correct sgRNA insertion. Sequence-verified plasmids were then transformed into *E. coli* Epi300 + pTA-mob for conjugation experiments, or isolated via column purification (Monarch Spin Plasmid Miniprep Kit, New England Biolabs) from *E. coli* and eluted with 40 *µ*L ddH_2_O for electroporation experiments.

### DNA delivery into *P. tricornutum*

Conjugation experiments were performed as previously described^7,8^, using *E. coli* Epi300 containing the conjugative helper plasmid pTA-mob and the plasmid to be delivered as a donor strain. Electroporation experiments were performed as described by Walker and colleagues^9^ with some modifications. Briefly,

*P. tricornutum* was adjusted to a concentration of 1 ·10^8^ cells/mL with a hemocytometer and plated onto 50% salt L1 1% agar solid media. Plates were then grown for 4-5 days, then scraped with 1.5 mL L1 into a 1.5 mL microfuge tube. Cells were then adjusted to 3-6·10^8^ cells/tube. Spheroplasting was performed with 10 *µ*L of alcalase (Sigma-Aldrich) diluted 10-fold in sterile ddH_2_O per tube, incubated for 20 minutes at room temperature on the lowest speed of a rotating mixer. After spheroplasting and washes, cells were resuspended to a final concentration of ∼ 1-3· 10^9^ cells/mL in ice-cold 385 mM D-sorbitol. 50 *µ*L of spheroplast preparation was mixed with 40 *µ*g of single-stranded salmon sperm DNA (Agilent) and 250 ng of prepared plasmid DNA (or 150 ng of positive control pPtGE27) in a microfuge tube, then transferred to pre-chilled electrocuvettes (VWR, 2 mm) and placed on ice. Electroporations were performed with a Bio-Rad Gene Pulser (Model No. 1652076) set to the following parameters: 500 V voltage, 25 *µ*F capacitance, and 400 Ω resistance. With these parameters, time constants of 7.8–8.8 milliseconds were typically seen. Immediately after electroporation the cuvette was placed back on ice, then 1 mL of L1 was gently pipetted on top of the reaction with tubes left to recover at room temperature for 10 minutes. After this brief recovery, the contents of the cuvettes were transferred to 50 mL centrifuge tubes containing 10 mL of L1, and left to recover under normal *P. tricornutum* growing conditions for 3 days. After 3 days of recovery, tubes were pelleted at 2k RCF at 18^*°*^C for 10 minutes, wherein supernatants were gently poured off and pellets were resuspended in 1–2 mL L1. 100 *µ*L of this was plated onto appropriate selective solid media conditions. For the experiments determining better conditions for selection with zeocin, the plasmid pPtGE27 was used, which contains both a *Sh ble* resistance gene (for zeocin selection) and nourseothricin (NTC) N-acetyl transferase (*nat)* gene (for NTC selection). Multi-day recoveries were tested by gently vortexing the recovery in the 50 mL centrifuge tube and pouring off ∼3.5 mL into a fresh 15 mL centrifuge tube on each time point tested. This portion of the recovery was then pelleted and resuspended as before, and plated out onto either 25% salt L1 solid media with 100 *µ*g/mL NTC, or zeocin-containing solid media as indicated. Efficiency of recovery with zeocin was reported as the ratio of transformants on zeocin-containing plates to NTC-containing plates, which was then compared to the recovery efficiency seen for conjugation experiments using the typical 25% salt L1 solid media with 100 *µ*g/mL zeocin or 100 *µ*g/mL NTC.

### Northern blot to detect sgRNA processing

A culture that had received the pPHYCUT plasmid with the *HIS, UMPS*, and *APT* sgRNAs via conjugation was grown in 50% salt L1 with 100 *µ*g/mL zeocin to a density of 2 · 10^6^ cells/mL. 50 mL of culture was harvested and pelleted at 6k RCF at 4^*°*^C, wherein the supernatant was gently poured off and the pellet was resuspended in 200 *µ*L TE buffer. This was then pipetted drop-wise into liquid nitrogen and ground into powder with a pre-chilled mortar and pestle. RNA extraction was performed with the Monarch™ Total RNA Miniprep Kit (New England Biolabs) according to manufacturer’s instructions. Purified RNA was run on a 1% agarose gel to confirm integrity. For the Northern blot, 20 *µ*L of the purified total RNA was mixed with 20 *µ*L of 2x Formamide dye (95% deionized formamide, 5 mM EDTA, 0.025% (w/v) bromophenol blue, 0.025% (w/v) xylene cyanol), while 0.05 fmols (5 *µ*L) of the ssDNA control (*HIS*-sgRNA, Supplementary table S9) was mixed with 5 *µ*L of 2x Formamide dye. Samples were heated at 95^*°*^C for 3 minutes before being loaded on a 10% denaturing acrylamide TBE gel (8 M urea, 1.5 mm Mini-PROTEAN® gel, Bio-Rad), along with 10 *µ*L of prepared Low Range ssRNA Ladder (New England Biolabs). The gel was run in TBE buffer at 150 V for approximately 55 minutes, when the dye front had traveled ~75% of the gel. The gel lane containing the ladder was excised and stained with ethidium bromide, and the remaining portion was transferred to a nylon membrane with a capillary transfer system in 0.5x TBE buffer with an ice pack at 180 mA for 60 minutes. The membrane was rinsed with 0.5x TBE and air dried, then UV crosslinked. The membrane was pre-hybridized with pre-warmed ULTRAhyb-oligo (Invitrogen) buffer at 42^*°*^C for 30 minutes with gentle agitation in a glass tube. 5 nM of a biotylinated ss-RNA probe (DE7017) was then added for hybridization overnight at 42^*°*^C. Membrane was then washed 2x30 minutes in stringent wash buffer (0.5% SDS (w/v), 300 mM NaCl, 30 mM sodium citrate, pH 7.0), followed by 2x5 minutes in wash buffer (1x PBS, 0.5% SDS (w/v)), then 2x5 minutes in blocking buffer (1x PBS, 0.5% SDS (w/v), 1% BSA (w/v)). The membrane was then incubated in blocking buffer for 30 minutes, then with streptavidin-alkaline phosphatase conjugate (Invitrogen, 1:10,000 in blocking buffer) for 30 minutes. The membrane was then washed 1x15 minutes in blocking buffer, 3x15 minutes in wash buffer, and 2x2 minutes in assay buffer (0.1 M diethanolamine, 1 mM MgCl_2_, pH 10.0). Chemiluminescent signal was then developed by incubating the membrane in 2 mL of Tropix CDP-Star solution for 5 minutes.

### Screening of targeted loci with PCRs and T7 endonuclease assays

After DNA delivery, *P. tricornutum* colonies on selective solid media were re-patched onto fresh selection plates for screening. The target loci were amplified with the Phire Plant Direct PCR Master Mix (ThermoScientific), using either a small amount of the streak (tip of a 10 *µ*L pipette) directly, or lysing a small amount of the streak in 30 *µ*L of TE at 98^*°*^C for 10 minutes and using 1 *µ*L of this as template. PCR reactions were carried out according to manufacturer’s instructions using 35 cycles, with a 2-step cycling protocol used for short (¡1 kb) amplicons and a 3-step cycling protocol with an annealing temperature of 65^*°*^C for longer amplicons. T7 endonuclease (T7E1) assays were performed on ∼ 500-900 bp amplicons to detect the presence of small indels. 2.5 *µ*L of sample amplicon was mixed with 2.5 *µ*L of wildtype amplicon to ensure homozygous mutations were still detected. The amplicon mixtures were heated at 95^*°*^C for 5 minutes then cooled at a rate of [0.2^*°*^C/s] to 50^*°*^C, wherein they were placed at -20^*°*^C for approximately 20 minutes. 0.2 *µ*L of T7E1 (New England Biolabs), 1.5 *µ*L of NEB2 buffer, and 8.3 *µ*L of ddH_2_O were added to each sample and incubated for 30 minutes at 37^*°*^C. PCR reactions were cleaned up over a column before being sent for Sanger sequencing at the London Regional Genomics Centre. Diploid editing events were deconvoluted with DECODR^58^. A list of all primers used for screening and sequencing is included in Supplementary table S9.

### Glycosyltransferase homolog search

Protein sequences for all enzymes in the KEGG pathways 00510 (*N*-glycan biosynthesis) and 00513 (Various types of *N*-glycan biosynthesis) were fetched for the organisms *Homo sapiens, Mus musculus, Arabidopsis thaliana, Glycine max, Zea mays, Saccharomyces cerevisiae, Danio rerio, Drosophila melanogaster*, and *Caenorhabditis elegans* using a Python script. BLASTp was run locally against the *P. tricornutum* Phatr3 proteome using the *N*-glycosylation pathway orthologs as queries with an expect value cut off of 1· 10^*−*^10. Putative *P. tricornutum* orthologs identified from this search were additionally verified with InterproScan^59^ to find conserved domains and structures. Transmembrane domains were predicted by TMHMM 2.0^60^ and DeepTMHMM^61^, signal peptides were predicted by signalP6.0^62^, and protein localization was predicted by DeepLoc2.1^63^. All glycosyltransferases identified are in Supplementary table S1.

## RNAseq experiment and data analysis

Five replicates of wildtype *P. tricornutum* were grown in 50 mL 50% salt L1 for approximately 4 weeks to a density of ∼4·10^7^ cells/mL, after which 2 ·10^8^ cells were harvested from each for RNA-extraction (pre-dilution samples). These cultures were then diluted to ∼1· 10^6^ cells/mL in fresh 50% salt L1 for a final volume of 300 mL culture. These 5 replicates were grown over a course of 17 days, with 2· 10^8^ cells harvested from each culture for RNA-extraction on days 4, 10, and 17. Culture density was monitored on sampling days via absorbance at 670 nm. RNA extractions were performed as described in the Northern blot section. RNAseq was performed at the London Regional Genomics Centre using the Illumina NextSeq 75 single end high output across five sequencing runs. Reads were trimmed with Trimmomatic^64^ version 0.36 with options LEADING:10 TRAILING:10. Processed reads were mapped to the ASM15095v2 reference genome including all unmapped scaffolds using Hisat2^65^ version 2.2.1. FeautureCounts^66^ was used to count the number of reads mapping to each annotated coding sequence feature in the Phatr3 annotation set. Counts tables across the 5 Illumina runs were merged and reads per kilobase (RPK) of each gene were calculated. All RPKs per sample were then summed and divided by 1 ·10^6^ to get the per million scaling factor for each sample. Then, each gene’s RPK was divided by the sample’s scaling factor to obtain its transcripts per kilobase million (TPM). For the *N*-glycosylation pathway heatmap, the TPM values for the 5 sample replicates for each time point were averaged. The *FucT* boxplots were overplotted with each individual replicate using a binwidth of 0.5. For the *PtFucT* read coverage histograms, reads mapped onto the genome for each Illumina run were merged and Samtools^67^ depth command was used to find the read counts mapped to each position of the region in each sample. The mean read count mapped per position across each replicate was calculated, and the total number of reads mapped per 5 nt bin was plotted on a log2 scale. To-scale diagrams of the *PtFucT* genomic regions according to the Phatr2 or Phatr3 genome annotations are aligned with the genomic coordinates underneath the coverage histograms.

### FucT phylogenetic tree generation

*P. tricornutum* FucT orthologs were identified by querying each of PtFucT1, PtFucT2, and PtFucT3 protein sequences against successively more exclusive databases with BLASTp. First, top hits from the standard nucleotide database against all organisms were found, then against a database exclusive of raphid pennate diatoms, then exclusive of all diatoms, and finally exclusive of all stramenopiles. For PtFucT1, whose top hits from an unrestricted database source were already mostly crustaceans and only one diatom species, databases searched against were all organisms exclusive of crustacea, only diatoms, stramenopiles exclusive of diatoms, and the SAR clade exclusive of stramenopiles. Structural predictions generated by AlphaFold 3^68^ for the orthologs identified from these searches, as well as characterized GT10 family FucTs from the CAZy database^69^, were used to identify the globular regions of the proteins. A multiple sequence alignment of the globular domains was generated with MAFFT^70^, and a maximum likelihood phylogenetic tree was generated from this alignment using IQtree3.0^71^ with 1000 Ultrafast bootstrap approximation replicates^72^. This tree was then visualized using Interactive Tree Of Life^73^. For all sequences used in this analysis, see Supplemental table S2.

### FucT mutant generation and screening

The pPHYCUT vector was cloned with sgRNAs targeting each of the three *FucT* genes in *P. tricornutum*, then conjugated into *P. tricornutum* via *E. coli*. Exconjugants were selected for on 25% salt L1 solid media containing 100 *µ*g/mL zeocin. Streaked out exconjugants were screened for editing at the targeted loci with primers surrounding each target site (Supplementary table S9) and T7E1 assays. Large deletions were screened by amplifying larger regions (∼3-4 kb) surrounding target sites. Four exconjugants that showed evidence of editing at all three targeted alleles were diluted and replated to obtain subclones. Subclones derived from each of the four exconjugants were then screened as before. Several subclones showing evidence of different editing patterns were selected for sequencing and further phenotypic characterization. The edits of the *FucT* genes were determined via Sanger sequencing of amplified target regions, and deconvolution of alleles was performed with DECODR. For *FucT* mutants generated by the non-multiplexing construct, single sgRNAs were cloned into pSS29 and conjugated into *P. tricornutum*, then screened as above.

### *α*(1,3)-fucosylation detection and phenotypic characterization of FucT mutants

FucT mutants and wildtype strains of *P. tricornutum* were seeded at a density of 1 ·10^6^ cells/mL to a volume of 15 mL in 50 mL centrifuge tubes, in 50% salt modified L1 media containing 4.4 mM urea as the sole nitrogen source and 0.5% glycerol (w/v). Each strain was seeded in triplicate. For each measurement, samples were read with four technical replicates in a 384-well plate (ThermoScientific, MaxiSorp clear 384 well plate). Every two days growth was monitored via absorbance readings at 670 nm, and chlorophyll autofluorescence was monitored via excitation at 440 nm and emission at 685 nm, using the BioTek Synergy H1 platereader. On days where extracellular medium was sampled for ELISA readings (days 4, 8, and 12), culture tubes were pelleted at 4k RCF for 10 minutes at 4^*°*^C and 300 *µ*L of supernatant sample were taken for analysis. Cultures were then briefly vortexed to resuspend. For protein quantification in the supernatant, a modified Bradford microassay (Bio-Rad) was used. Briefly, standards of Bovine Serum Albumin were made ranging in concentration from 0-25 ng/*µ*L, using the experimental media as diluent. 25 *µ*L of each supernatant sample or standard was pipetted in four replicates, and 40 *µ*L of Bradford reagent (Bio-Rad) was added to each well. Absorbance at 595 nm was taken after a 5 minute incubation. A modified version of the Abcam indirect ELISA protocol was used to quantify the levels of *α*(1,3)-fucosylation in samples. Briefly, standards of Horseradish peroxidase (HRP, Sigma-Aldrich) were made ranging in concentration from 0-40 ng/mL, using the experimental media as diluent. 15 *µ*L of each supernatant sample or standard was pipetted in four replicates and incubated covered at room temperature for 2 hours. The samples were then washed with 3x50 *µ*L PBS, then incubated in 50 *µ*L carbo-free blocking buffer (Vector Laboratories) at 4^*°*^C overnight. The next day, the samples were washed again with 3x50 *µ*L PBS, then incubated in 25 *µ*L of primary antibody solution (Rabbit anti-*α*(1,3)-fucose antibody from Agrisera, diluted 1:6,000 in PBS) for 2 hours at room temperature. Samples were then washed with 4x50 *µ*L of PBS, then incubated in secondary antibody solution (Cytiva Amersham ECL Rabbit IgG, HRP-linked whole Ab (from donkey), diluted 1:8,000 in PBST) for 1.5 hours at room temperature. The samples were washed with 4x50 *µ*L of PBST, then 25 *µ*L of TMB/E reagent (Millipore) was added to each well. Colour was allowed to develop for approximately 15 minutes at room temperature, after which 25 *µ*L of 0.1 M NaF stop solution was added to each well, and absorbance readings were taken at 650 nm.

### ELISA experiment data analyses

All technical replicate data was filtered to remove outlier replicates where the difference between the data point and the median is greater than 1.5 times the interquartile range, leaving at least 3 technical replicates for each sample. Technical replicate data was then averaged for each biological replicate. For growth curves, absorbance at 670 nm was blanked by the experimental media. Protein concentrations of supernatant samples were determined from the standard curve generated by the Bradford assay for each sampling day. Fucose reactivity was determined relative to the HRP standard curve generated on each sampling day. Fucose reactivity for each sample was normalized by either the culture density as determined by absorbance at 670 nm, or protein concentration of the supernatant sample. The fold change of fucose reactivity was determined by dividing the normalized fucose reactivity of the mutant strain by the average normalized fucose reactivity of the three wildtype strains (9 biological replicates total), wherein the log2 fold change was plotted relative to wildtype. All p-values were determined by comparison of the mutant strain to the wildtype average on each day with an unpaired students T-test, then corrected with a Benjamini-Hochberg adjustment.

## Acknowledgments

We thank members of the Karas Lab for comments on the manuscript.

## Data availability

The RNAseq datasets have been deposited in the Short Read Archive with the accession number PRJNA1348894. The PHYCUT plasmids have been deposited in AddGene.

## Author contributions

Conceptualization, E.E.S, S.S.S., L.S.G., G.B.G., D.R.E.; methodology, E.E.S., L.S.G., K.H.D., T.S.B.; investigation, E.S.S., L.S.G. S.S.S., K.H.D.; writing, E.S.S., G.B.G., D.R.E.; funding acqusition, D.R.E., G.B.G.; supervision, D.R.E., G.B.G.

## Declaration of interests

The authors declare no competing interests.

**Supplemental information index**

**Supplemental Figure S1** Optimization of conditions used to select for *P. tricornutum* electrotransformants.

**Supplemental Figure S2** Screening P. tricornutum expressing APT -targeting guides in the PHYCUT contruct.

**Supplemental Figure S3** Phylogenetic analysis of the three GT10-family PtFucTs.

**Supplemental Figure S4** FucT PHYCUT exconjugants screened for the presence of editing at targeted loci.

**Supplemental Figure S5** FucT PHYCUT subclones screened for the presence of editing at targeted loci.

**Supplemental Figure S6** FucT PHYCUT subclones screened for defined-length deletions between adjacent target sites.

**Supplemental Figure S7** Indirect ELISA protocol allows sensitive levels of fucosylation detection.

**Supplemental Figure S8** Culture density-normalized fucosylation of secreted proteins relative to wild type.

**Supplemental Figure S9** Protein-normalized fucosylation of secreted proteins relative to wild type.

**Supplemental Figure S10** Growth curves for all strains studied in ELISA experiment.

**Supplemental Table S1** *N*-glycosylation pathway orthologs identified in *P. tricornutum*.

**Supplemental Table S2** FucT homologs used to construct phylogenetic tree for the PtFucTs.

**Supplemental Table S3** The three FucT genes predicted in *P. tricornutum* and their annotated coding regions.

**Supplemental Table S4** *PtFucT* -targeted sgRNAs used in this study.

**Supplemental Table S5** Genetic edits made to the *PtFucT* genes for strains generated using the non-multiplexing Cas9 construct.

**Supplemental Table S6** Genetic edits made to the *PtFucT* genes for strains generated using the PHYCUT construct.

**Supplemental Table S7** Plasmids used in this study.

**Supplemental Table S8** Genetic parts used to create constructs for this study. For sgRNA inserts, BsaI recognition sites are shown in red text, the 4 nt overhangs generated by BsaI digestion are shown in blue text, sgRNA spacer sequences are bolded, and Csy4 recognition sites are underlined.

**Supplemental Table S9** DNA oligonucleotides used in this study. For sgRNA inserts, BsaI recognition sites are shown in red text, the 4 nt overhangs generated by BsaI digestion are shown in blue text, sgRNA spacer sequences are bolded, and Csy4 recognition sites are underlined.

## References

1. Nelson, D. M., Tréguer, P., Brzezinski, M. A., Leynaert, A., and Quéguiner, B. (1995). Production and dissolution of biogenic silica in the ocean: revised global estimates, comparison with regional data and relationship to biogenic sedimentation. Global Biogeochemical Cycles 9, 359–372.

2. Bowler, C., Vardi, A., and Allen, A. E. (2010). Oceanographic and biogeochemical insights from diatom genomes. Annual Review of Marine Science 2, 333–365.

3. Bowler, C., Allen, A. E., Badger, J. H., Grimwood, J., Jabbari, K., Kuo, A., Maheswari, U., Martens, C., Maumus, F., and Otillar, R. P. (2008). The phaeodactylum genome reveals the evolutionary history of diatom genomes. Nature 456, 239.

4. Giguere, D. J., Bahcheli, A. T., Slattery, S. S., Patel, R. R., Browne, T. S., Flatley, M., Karas, B. J., Edgell, D. R., and Gloor, G. B. (2022). Telomere-to-telomere genome assembly of Phaeodactylum tricornutum. PeerJ 10, e13607.

5. Slattery, S. S., Diamond, A., Wang, H., Therrien, J. A., Lant, J. T., Jazey, T., Lee, K., Klassen, Z., Desgagné-Penix, I., Karas, B. J., and Edgell, D. R. (2018). An expanded plasmid-based genetic tool-box enables Cas9 genome editing and stable maintenance of synthetic pathways in Phaeodactylum tricornutum. ACS Synthetic Biology 7, 328–338.

6. Chu, L., Ewe, D., Bártulos, C. R., Kroth, P. G., and Gruber, A. (2016). Rapid induction of GFP expression by the nitrate reductase promoter in the diatom Phaeodactylum tricornutum. PeerJ 4, e2344.

7. Slattery, S. S., Giguere, D. J., Stuckless, E. E., Shrestha, A., Briere, L.-A. K., Galbraith, A., Reaume, S., Boyko, X., Say, H. H., Browne, T. S. et al. (2022). Phosphate-regulated expression of the sars-cov-2 receptor-binding domain in the diatom phaeodactylum tricornutum for pandemic diagnostics. Scientific Reports 12, 7010.

8. Karas, B. J., Diner, R. E., Lefebvre, S. C., McQuaid, J., Phillips, A. P., Noddings, C. M., Brunson, J. K., Valas, R. E., Deerinck, T. J., Jablanovic, J. et al. (2015). Designer diatom episomes delivered by bacterial conjugation. Nature Communications 6, 6925.

9. Walker, E. J. L., Pampuch, M., Tran, G., and Karas, B. J. (2024). Spheroplasted cells: a game changer for DNA delivery to diatoms. bioRxiv (2010–2024).

10. Apt, K. E., Grossman, A. R., and Kroth-Pancic, P. G. (1996). Stable nuclear transformation of the diatom Phaeodactylum tricornutum. Molecular and General Genetics MGG 252, 572–579.

11. Nymark, M., Sharma, A. K., Hafskjold, M. C. G., Sparstad, T., Bones, A. M., and Winge, P. (2017). CRISPR/Cas9 gene editing in the marine diatom Phaeodactylum tricornutum. Bio-protocol 7, e2442–e2442.

12. Stukenberg, D., Zauner, S., Dell’Aquila, G., and Maier, U. G. (2018). Optimizing CRISPR/Cas9 for the diatom Phaeodactylum tricornutum. Frontiers in Plant Science 9, 740.

13. Serif, M., Dubois, G., Finoux, A.-L., Teste, M.-A., Jallet, D., and Daboussi, F. (2018). One-step generation of multiple gene knock-outs in the diatom Phaeodactylum tricornutum by DNA-free genome editing. Nature Communications 9, 3924.

14. Novoveská, L., Ross, M. E., Stanley, M. S., Pradelles, R., Wasiolek, V., and Sassi, J.-F. (2019). Microalgal carotenoids: A review of production, current markets, regulations, and future direction. Marine Drugs 17, 640.

15. Gilmour, D. J. (2019). Microalgae for biofuel production. Advances in Applied Microbiology 109, 1–30.

16. Oey, M., Sawyer, A. L., Ross, I. L., and Hankamer, B. (2016). Challenges and opportunities for hydrogen production from microalgae. Plant Biotechnology Journal 14, 1487–1499.

17. Barolo, L., Abbriano, R. M., Commault, A. S., George, J., Kahlke, T., Fabris, M., Padula, M. P., Lopez, A., Ralph, P. J., and Pernice, M. (2020). Perspectives for glyco-engineering of recombinant biopharmaceuticals from microalgae. Cells 9, 633.

18. Dehghani, J., Adibkia, K., Movafeghi, A., Maleki-Kakelar, H., Saeedi, N., and Omidi, Y. (2020). Towards a new avenue for producing therapeutic proteins: Microalgae as a tempting green biofactory. Biotechnology Advances 40, 107499.

19. Hempel, F., Lau, J., Klingl, A., and Maier, U. G. (2011). Algae as protein factories: expression of a human antibody and the respective antigen in the diatom Phaeodactylum tricornutum. PloS one 6, e28424.

20. Rosales-Mendoza, S., Solís-Andrade, K. I., Márquez-Escobar, V. A., González-Ortega, O., and Bañuelos-Hernandez, B. (2020). Current advances in the algae-made biopharmaceuticals field. Expert Opinion on Biological Therapy 20, 751–766.

21. Reichert, J. M. Metrics for antibody therapeutics development. vol. 2. Taylor Francis. ISBN 1942-0862 (2010):(695–700).

22. Baïet, B., Burel, C., Saint-Jean, B., Louvet, R., Menu-Bouaouiche, L., Kiefer-Meyer, M.-C., Mathieu-Rivet, E., Lefebvre, T., Castel, H., Carlier, A. et al. (2011). N-glycans of Phaeodactylum tricornutum diatom and functional characterization of its N-acetylglucosaminyltransferase I enzyme. Journal of Biological Chemistry 286, 6152–6164.

23. Mathieu-Rivet, E., Kiefer-Meyer, M.-C., Vanier, G., Ovide, C., Burel, C., Lerouge, P., and Bardor, M. (2014). Protein N-glycosylation in eukaryotic microalgae and its impact on the production of nuclear expressed biopharmaceuticals. Frontiers in Plant Science 5, 359.

24. Dumontier, R., Loutelier-Bourhis, C., Walet-Balieu, M.-L., Burel, C., Mareck, A., Afonso, C., Lerouge, P., and Bardor, M. (2021). Identification of N-glycan oligomannoside isomers in the diatom Phaeodactylum tricornutum. Carbohydrate Polymers 259, 117660.

25. van Bockstaele-Fuentes, J., Mati-Baouche, N., Lupette, J., Gargouch, N., Rivet, E., Lerouge, P., and Bardor, M. (2025). An overview of protein N-glycosylation diversity in microalgae. Frontiers in Plant Science 16, 1669918.

26. Van Beers, M. M., and Bardor, M. (2012). Minimizing immunogenicity of biopharmaceuticals by controlling critical quality attributes of proteins. Biotechnology journal 7, 1473–1484.

27. Manduzio, H., Fitchette, A.-C., Hrabina, M., Chabre, H., Batard, T., Nony, E., Faye, L., Moingeon, P., and Gomord, V. (2012). Glycoproteins are species-specific markers and major ige reactants in grass pollens. Plant biotechnology journal 10, 184–194.

28. Haurwitz, R. E., Jinek, M., Wiedenheft, B., Zhou, K., and Doudna, J. A. (2010). Sequence-and structure-specific RNA processing by a CRISPR endonuclease. Science 329, 1355–1358.

29. Kurata, M., Wolf, N. K., Lahr, W. S., Weg, M. T., Kluesner, M. G., Lee, S., Hui, K., Shiraiwa, M., Webber, B. R., and Moriarity, B. S. (2018). Highly multiplexed genome engineering using CRISPR/Cas9 gRNA arrays. PloS one 13, e0198714.

30. McCarty, N. S., Graham, A. E., Studená, L., and Ledesma-Amaro, R. (2020). Multiplexed CRISPR technologies for gene editing and transcriptional regulation. Nature Communications 11, 1281.

31. Chu, L., Ewe, D., Bártulos, C. R., Kroth, P. G., and Gruber, A. (2016). Rapid induction of GFP expression by the nitrate reductase promoter in the diatom Phaeodactylum tricornutum. PeerJ 4, e2344.

32. Adler-Agnon, Z., Leu, S., Zarka, A., Boussiba, S., and Khozin-Goldberg, I. (2018). Novel promoters for constitutive and inducible expression of transgenes in the diatom Phaeodactylum tricornutum under varied nitrate availability. Journal of Applied Phycology 30, 2763–2772.

33. Slattery, S. S., Wang, H., Giguere, D. J., Kocsis, C., Urquhart, B. L., Karas, B. J., and Edgell, D. R. (2020). Plasmid-based complementation of large deletions in Phaeodactylum tricornutum biosynthetic genes generated by Cas9 editing. Scientific Reports 10, 13879.

34. Zhang, Y., Ge, X., Yang, F., Zhang, L., Zheng, J., Tan, X., Jin, Z.-B., Qu, J., and Gu, F. (2014). Comparison of non-canonical PAMs for CRISPR/Cas9-mediated dna cleavage in human cells. Scientific Reports 4, 5405.

35. Diamond, A., Diaz-Garza, A. M., Li, J., Slattery, S. S., Merindol, N., Fantino, E., Meddeb-Mouelhi, F., Karas, B. J., Barnabé, S., and Desgagné-Penix, I. (2023). Instability of extrachromosomal DNA transformed into the diatom Phaeodactylum tricornutum. Algal Research 70, 102998.

36. Sakaguchi, T., Nakajima, K., and Matsuda, Y. (2011). Identification of the UMP synthase gene by establishment of uracil auxotrophic mutants and the phenotypic complementation system in the marine diatom Phaeodactylum tricornutum. Plant Physiology 156, 78–89.

37. Zhang, P., Burel, C., Plasson, C., Kiefer-Meyer, M.-C., Ovide, C., Gügi, B., Wan, C., Teo, G., Mak, A., Song, Z. et al. (2019). Characterization of a GDP-fucose transporter and a fucosyltransferase involved in the fucosylation of glycoproteins in the diatom Phaeodactylum tricornutum. Frontiers in Plant Science 10, 610.

38. Both, P., Sobczak, L., Breton, C., Hann, S., Nöbauer, K., Paschinger, K., Kozmon, S., Mucha, J., and Wilson, I. B. (2011). Distantly related plant and nematode core α1, 3-fucosyltransferases display similar trends in structure–function relationships. Glycobiology 21, 1401–1415.

39. Xie, X., Yang, J., Du, H., Chen, J., Sanganyado, E., Gong, Y., Du, H., Chen, W., Liu, Z., and Liu, X. (2023). Golgi fucosyltransferase 1 reveals its important role in α-1, 4-fucose modification of n-glycan in crispr/cas9 diatom phaeodactylum tricornutum. Microbial Cell Factories 22, 6.

40. Bardor, M., Balieu, J., Perruchon, O., Loutelier-Bourhis, C., Afonso, C., Mathieu-Rivet, E., and Lerouge, P. (2024). Maturation of protein N-glycans depends on phaeodactylum tricornutum ecotypes and results in the synthesis of novel complex n-glycans in pt3. Available at SSRN 4912887.

41. Rastogi, A., Maheswari, U., Dorrell, R. G., Vieira, F. R. J., Maumus, F., Kustka, A., McCarthy, J., Allen, A. E., Kersey, P., Bowler, C. et al. (2018). Integrative analysis of large scale transcriptome data draws a comprehensive landscape of Phaeodactylum tricornutum genome and evolutionary origin of diatoms. Scientific Reports 8, 4834.

42. Deschamps, P., and Moreira, D. (2012). Reevaluating the green contribution to diatom genomes. Genome Biology and Evolution 4, 683–688.

43. Niazi, S. K. (2025). Continuous manufacturing of recombinant drugs: Comprehensive analysis of cost reduction strategies, regulatory pathways, and global implementation. Pharmaceuticals 18, 1157.

44. Qian, L., Lin, X., Gao, X., Khan, R. U., Liao, J.-Y., Du, S., Ge, J., Zeng, S., and Yao, S. Q. (2023). The dawn of a new era: targeting the “undruggables” with antibody-based therapeutics. Chemical reviews 123, 7782–7853.

45. El-Araby, R. (2024). Biofuel production: exploring renewable energy solutions for a greener future. Biotechnology for Biofuels and Bioproducts 17, 129.

46. Atnoorkar, S., Wiatrowski, M., Newes, E., Davis, R., and Peterson, S. O. (2024). Algae to hefa: Economics and potential development trajectories for deployment in the united states. Biofuels, Bioproducts and Biorefining (Online) 18.

47. Hao, X., Chen, W., Amato, A., Jouhet, J., Maréchal, E., Moog, D., Hu, H., Jin, H., You, L., and Huang, F. (2022). Multiplexed CRISPR/Cas9 editing of the long-chain acyl-CoA synthetase family in the diatom Phaeodactylum tricornutum reveals that mitochondrial ptACSL3 is involved in the synthesis of storage lipids. New Phytologist 233, 1797–1812.

48. Moosburner, M. A., Gholami, P., McCarthy, J. K., Tan, M., Bielinski, V. A., and Allen, A. E. (2020). Multiplexed knockouts in the model diatom Phaeodactylum by episomal delivery of a selectable Cas9. Frontiers in Microbiology 11, 5.

49. Taparia, Y., Dolui, A. K., Boussiba, S., and Khozin-Goldberg, I. (2022). Multiplexed genome editing via an RNA polymerase II promoter-driven sgRNA array in the diatom Phaeodactylum tricornutum: insights into the role of StLDP. Frontiers in Plant Science 12, 784780.

50. Sternberg, S. H., Haurwitz, R. E., and Doudna, J. A. (2012). Mechanism of substrate selection by a highly specific CRISPR endoribonuclease. RNA 18, 661–672.

51. Schramm, L., and Hernandez, N. (2002). Recruitment of RNA polymerase III to its target promoters. Genes & Development 16, 2593–2620.

52. Strasser, R., Altmann, F., Mach, L., Glössl, J., and Steinkellner, H. (2004). Generation of Arabidopsis thaliana plants with complex N-glycans lacking β1, 2-linked xylose and core α1, 3-linked fucose. FEBS Letters 561, 132–136.

53. Jansing, J., Sack, M., Augustine, S. M., Fischer, R., and Bortesi, L. (2019). CRISPR/Cas9-mediated knockout of six glycosyltransferase genes in Nicotiana benthamiana for the production of recombinant proteins lacking β-1, 2-xylose and core α-1, 3-fucose. Plant Biotechnology Journal 17, 350–361.

54. Boulogne, I., Toustou, C., and Bardor, M. (2025). Meta-analysis of RNA-Seq datasets allows a better understanding of P. tricornutum cellular biology, a requirement to improve the production of biologics. Scientific Reports 15, 3603.

55. Toustou, C., Boulogne, I., Gonzalez, A.-A., and Bardor, M. (2024). Comparative RNA-Seq of ten Phaeodactylum tricornutum accessions: unravelling criteria for robust strain selection from a bio-production point of view. Marine Drugs 22, 353.

56. Hempel, F., and Maier, U. G. (2012). An engineered diatom acting like a plasma cell secreting human IgG antibodies with high efficiency. Microbial Cell Factories 11, 126.

57. Marillonnet, S., and Grützner, R. (2020). Synthetic DNA assembly using golden gate cloning and the hierarchical modular cloning pipeline. Current Protocols in Molecular Biology 130, e115.

58. Bloh, K., Kanchana, R., Bialk, P., Banas, K., Zhang, Z., Yoo, B.-C., and Kmiec, E. B. (2021). Deconvolution of complex DNA repair (DECODR): establishing a novel deconvolution algorithm for comprehensive analysis of CRISPR-edited sanger sequencing data. The CRISPR Journal 4, 120–131.

59. Jones, P., Binns, D., Chang, H.-Y., Fraser, M., Li, W., McAnulla, C., McWilliam, H., Maslen, J., Mitchell, A., Nuka, G. et al. (2014). Interproscan 5: genome-scale protein function classification. Bioinformatics 30, 1236–1240.

60. Krogh, A., Larsson, B., Von Heijne, G., and Sonnhammer, E. L. (2001). Predicting transmembrane protein topology with a hidden Markov model: application to complete genomes. Journal of Molecular Biology 305, 567–580.

61. Hallgren, J., Tsirigos, K. D., Pedersen, M. D., Almagro Armenteros, J. J., Marcatili, P., Nielsen, H., Krogh, A., and Winther, O. (2022). Deeptmhmm predicts alpha and beta transmembrane proteins using deep neural networks. biorxiv (2022–04).

62. Teufel, F., Almagro Armenteros, J. J., Johansen, A. R., Gíslason, M. H., Pihl, S. I., Tsirigos, K. D., Winther, O., Brunak, S., von Heijne, G., and Nielsen, H. (2022). Signalp 6.0 predicts all five types of signal peptides using protein language models. Nature Biotechnology 40, 1023–1025.

63. Ødum, M. T., Teufel, F., Thumuluri, V., Almagro Armenteros, J. J., Johansen, A. R., Winther, O., and Nielsen, H. (2024). Deeploc 2.1: multi-label membrane protein type prediction using protein language models. Nucleic Acids Research 52, W215–W220.

64. Bolger, A. M., Lohse, M., and Usadel, B. (2014). Trimmomatic: a flexible trimmer for illumina sequence data. Bioinformatics 30, 2114–2120.

65. Kim, D., Paggi, J. M., Park, C., Bennett, C., and Salzberg, S. L. (2019). Graph-based genome alignment and genotyping with HISAT2 and HISAT-genotype. Nature Biotechnology 37, 907–915.

66. Liao, Y., Smyth, G. K., and Shi, W. (2014). featurecounts: an efficient general purpose program for assigning sequence reads to genomic features. Bioinformatics 30, 923–930.

67. Li, H., Handsaker, B., Wysoker, A., Fennell, T., Ruan, J., Homer, N., Marth, G., Abecasis, G., Durbin, R., and Subgroup, . G. P. D. P. (2009). The sequence alignment/map format and samtools. Bioinformatics 25, 2078–2079.

68. Abramson, J., Adler, J., Dunger, J., Evans, R., Green, T., Pritzel, A., Ronneberger, O., Willmore, L., Ballard, A. J., Bambrick, J. et al. (2024). Accurate structure prediction of biomolecular interactions with alphafold 3. Nature 630, 493–500.

69. Drula, E., Garron, M.-L., Dogan, S., Lombard, V., Henrissat, B., and Terrapon, N. (2022). The carbohydrate-active enzyme database: functions and literature. Nucleic Acids Research 50, D571–D577.

70. Katoh, K., Rozewicki, J., and Yamada, K. D. (2019). MAFFT online service: multiple sequence alignment, interactive sequence choice and visualization. Briefings in Bioinformatics 20, 1160–1166.

71. Wong, T. K., Ly-Trong, N., Ren, H., Baños, H., Roger, A. J., Susko, E., Bielow, C., De Maio, N., Goldman, N., Hahn, M. W. et al. (2025). Iq-tree 3: Phylogenomic inference software using complex evolutionary models.

72. Hoang, D. T., Chernomor, O., Von Haeseler, A., Minh, B. Q., and Vinh, L. S. (2018). UFBoot2: improving the ultrafast bootstrap approximation. Molecular Biology and Evolution 35, 518–522.

73. Letunic, I., and Bork, P. (2024). Interactive Tree of Life (iTOL) v6: recent updates to the phylogenetic tree display and annotation tool. Nucleic Acids Research 52, W78–W82.

