## Supplemental Figures for "PHYCUT: Scalable multiplex CRISPR/Cas9 editing for genome engineering in the diatom *Phaeodactylum tricornutum*"

### Supplemental Results

Proteins predicted to be orthologous to human enzymes were identified from the *P. tricornutum* genome, as described in the methods section. To confirm that the predicted glycosyltransferase enzymes were expressed in our strain, we used RNAseq to assess the transcripts per kilobase million (TPMs) at four different timepoints over 17 days of growth (Figure 5B). As with other eukaryotes, the process by which a lipid-linked oligosaccharide (LLO) precursor is built and then transferred to the acceptor asparagine residue of nascent polypeptides in the ER is mostly conserved. Homologs of ALG10 and MOGS, which respectively add and remove the third glucose (Glc) moiety onto the $GlcNAc_2Man_9Glc_3$ precursor in other eukaryotes, have not been identified. In addition, while two STT3 homologs were identified, no other OST complex subunits have been found. The STT3 subunit contains the catalytic active site of the OST and has been shown to be the only necessary OST subunit required for *N*-glycan transfer from the LLO to polypeptide, with other subunits determining donor-substrate specificity[1,2]. Predicted *P. tricornutum* glycosyltransferases were all found to be expressed to some degree, but we noted that transcript levels for ALG8 (Phatr3_J44905), which adds the second Glc moiety to the LLO precursor, were quite low for 3 of the 4 time points (TPM$<$1). Taken together, this may explain *P. tricornutum*'s likely $GlcNAc_2Man_9Glc_2$-LLO donor, or potentially even $GlcNAc_1Man_9Glc_1$-LLO donor, due to relaxed OST specificity.

Upon peptide transfer and deglucosylation of the *N*-glycan into Man9, the *N*-glycoprotein is transported to the Golgi apparatus and processed by $\alpha$-mannosidases MAN1B (Phatr3_J52346) and MAN1A (Phatr3_EG02374) to varying degrees into the oligomannosides Man5-Man9, which have been characterized in *P. tricornutum* to be the same as those produced by mammals[3]. Importantly, *P. tricornutum* encodes for *N*-acetylglucosaminyltransferase (MGAT1 or GnT I, Phatr3_J54844), which is responsible for the first step in the formation of complex *N*-glycans and was found to rescue complex *N*-glycan formation in a MGAT1-deficient CHO cell line[4]. In our search of glycosyltransferase homologs, we identified a dual-purpose MAN2-MGAT2 enzyme in *P. tricornutum* (Phatr3_J52248), which was also recently discussed by Bardor and colleagues[5]. The presence of these two MGAT enzymes, which add a GlcNAc residue to each arm of the Man5 core, indicates that *P. tricornutum* is capable of making complex *N*-glycans and would act as an excellent starting point for further humanization. However, experimental evidence of a second GlcNAc residue being added has not been seen yet, and levels of complex or hybrid *N*-glycans reported by mass spectrometry are low, with the majority of *N*-glycoproteins made by *P. tricornutum* containing oligomannosides[4,6,7]. Notably, the MGAT1 homolog is expressed at a low level, which may explain why complex *N*-glycans are seen at relatively low levels in *P. tricornutum*, with oligomannose glycans being the predominant species. Relatively high levels of HEXO1/3 expression, which are thought to degrade the terminal GlcNAc residue(s) from complex *N*-glycans in *P. tricornutum*, may also contribute to why such low levels of terminal GlcNAc-containing glycans are seen. It is after the actions of MGAT1 and potentially MGAT2 that the pathway of complex *N*-glycosylation in *P. tricornutum* diverges from that of humans. *P. tricornutum* encodes an ortholog of XylT (Phatr3_J45496), which adds a $\beta$(1,2)-xylose to the $\beta$(1,4)-mannose in plants and was recently discovered to also decorate the penultimate GlcNAc of the chitobiose core in the *P.tricornutum* ecotype Pt3[5], and B4GALT (Phatr3_J46140), predicted to add $\beta$(1,4)-galactose moieties to the terminal *N*-GlcNAc residues of complex *N*-glycan structures. Of relevance, after the action of MGAT1, MAN2, and potentially MGAT2, the glycan may be fucosylated by the three FucT genes identified in the genome.

### References

1. Kelleher, D. J., and Gilmore, R. (2006). An evolving view of the eukaryotic oligosaccharyltransferase. Glycobiology 16, 47R–62R.

2. Castro, O., Movsichoff, F., and Parodi, A. J. (2006). Preferential transfer of the complete glycan is

determined by the oligosaccharyltransferase complex and not by the catalytic subunit. Proceedings of the National Academy of Sciences 103, 14756–14760.

3. Dumontier, R., Loutelier-Bourhis, C., Walet-Balieu, M.-L., Burel, C., Mareck, A., Afonso, C., Lerouge, P., and Bardor, M. (2021). Identification of N-glycan oligomannoside isomers in the diatom *Phaeodactylum tricornutum*. Carbohydrate Polymers 259, 117660.

4. Baïet, B., Burel, C., Saint-Jean, B., Louvet, R., Menu-Bouaouiche, L., Kiefer-Meyer, M.-C., Mathieu-Rivet, E., Lefebvre, T., Castel, H., Carlier, A. et al. (2011). N-glycans of *Phaeodactylum tricornutum* diatom and functional characterization of its N-acetylglucosaminyltransferase I enzyme. Journal of Biological Chemistry 286, 6152–6164.

5. Bardor, M., Balieu, J., Perruchon, O., Loutelier-Bourhis, C., Afonso, C., Mathieu-Rivet, E., and Lerouge, P. (2024). Maturation of protein N-glycans depends on phaeodactylum tricornutum ecotypes and results in the synthesis of novel complex n-glycans in pt3. Available at SSRN 4912887.

6. Xie, X., Du, H., Chen, J., Aslam, M., Wang, W., Chen, W., Li, P., Du, H., and Liu, X. (2021). Global profiling of N-glycoproteins and N-glycans in the diatom *Phaeodactylum tricornutum*. Frontiers in Plant Science 12, 779307.

7. Vanier, G., Hempel, F., Chan, P., Rodamer, M., Vaudry, D., Maier, U. G., Lerouge, P., and Bardor, M. (2015). Biochemical characterization of human anti-hepatitis b monoclonal antibody produced in the microalgae phaeodactylum tricornutum. PLoS One 10, e0139282.

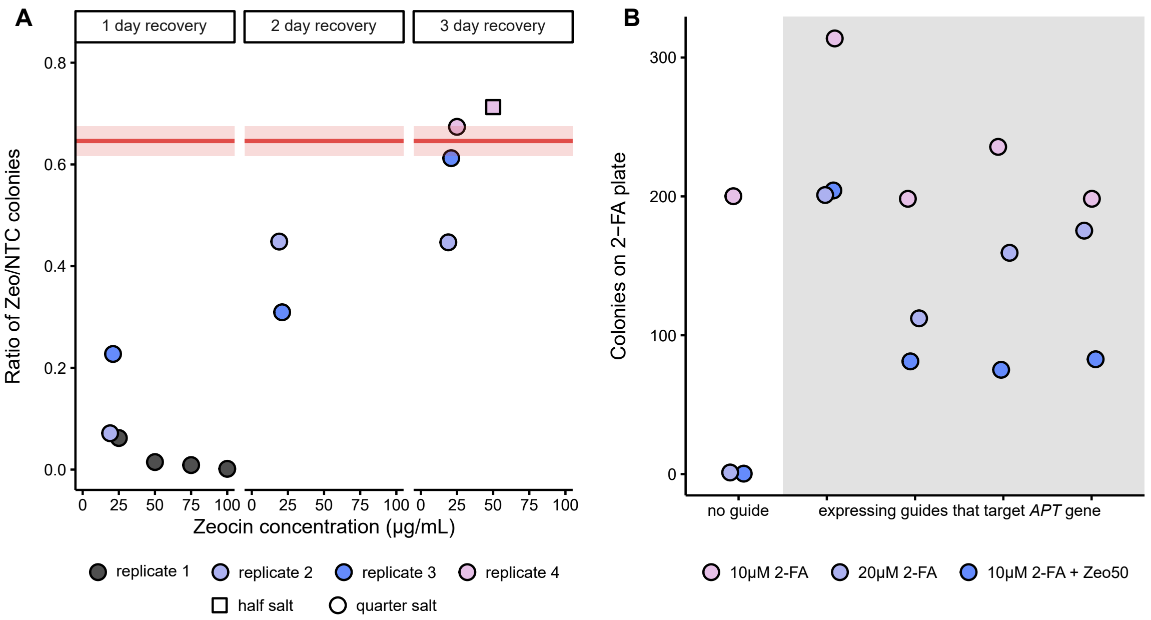


**Figure S 1.** Optimization of conditions used to select for *P. tricornutum* electrotransformants. A) The ratio of pPtGE27 transformants seen on zeocin-containing plates to transformants on 25% salt L1 + 100 $\mu$g/mL solid media. Increased recovery times, decreased antibiotic concentration, and increased salt concentration were tested to improve zeocin selection recovery. The solid and translucent red lines represent the average and standard deviation, respectively, of the ratio of pPtGE27 exconjugants on 25% salt L1 + 100 $\mu$g/mL zeocin to 25% salt L1 + 100 $\mu$g/mL NTC solid media (n=3). B) Increasing 2-FA concentration from 10 $\mu$M to 20 $\mu$M, or combining 10 $\mu$M 2-FA with 50 $\mu$g/mL zeocin, eliminates the small background colonies seen with 10 $\mu$M 2-FA selection only.

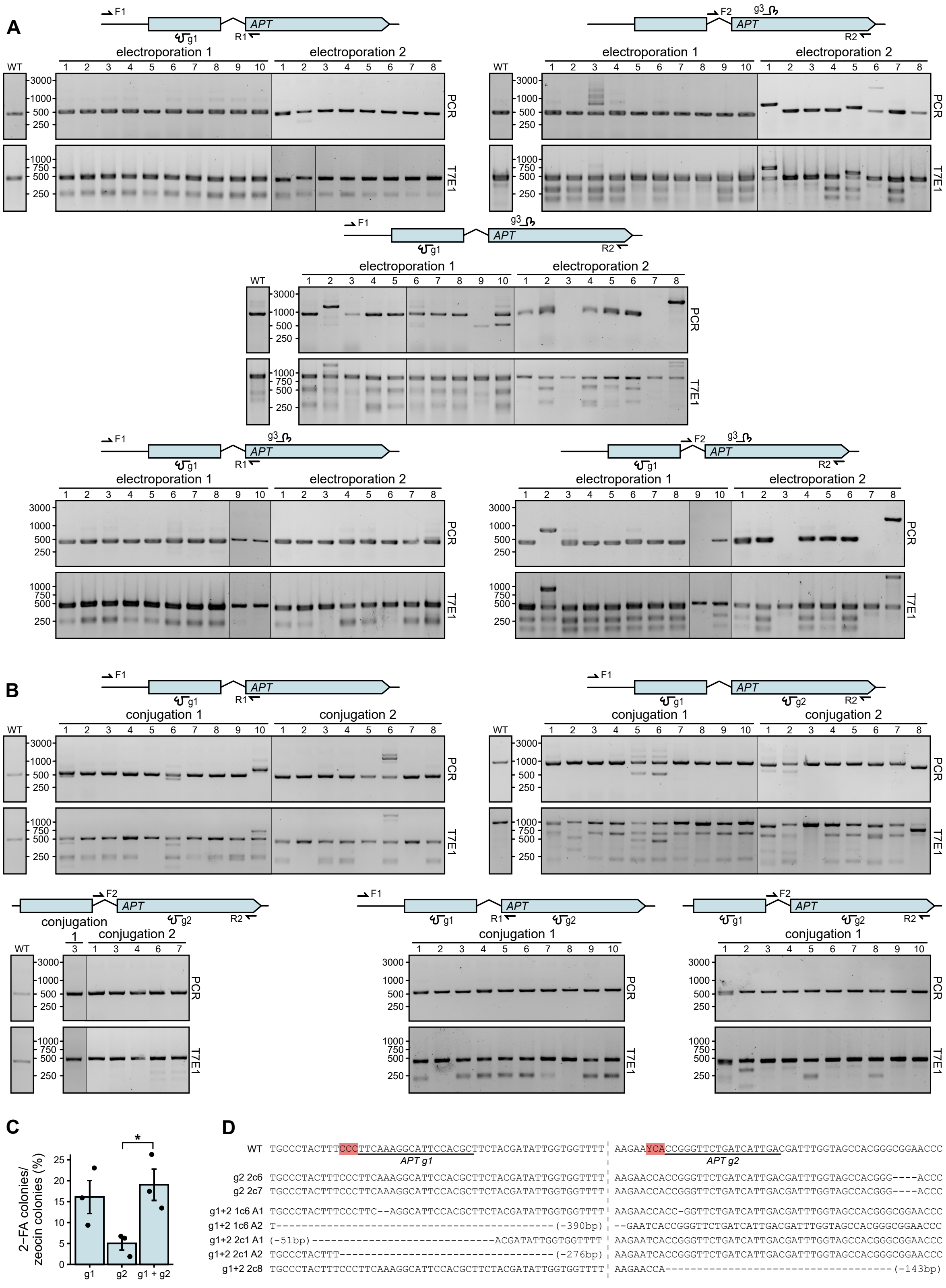


**Figure S 2.** Legend on following page

**Figure S 2.** Screening *P. tricornutum* expressing *APT*-targeting guides in the PHYCUT contruct. A) *APT*-sgRNA-g1 and g3 were cloned individually and together into the PHYCUT plasmid and electroporated into *P. tricornutum*. *APT* knock outs were selected for on 50% salt L1 solid media with 10 $\mu$M 2-FA and 50 $\mu$g/mL zeocin. The guides and screening primers used are indicated by the diagram above each set of agarose gels of PCR products (top) and T7E1 assays (bottom). Sizes (in bp) are indicated next to the gel images. Black horizontal lines separate colonies derived from two independent electroporations. WT indicates PCR amplicons and T7E1 digests from untreated *P. tricornutum*. T7E1 digests contain a mixture of PCR amplicons from WT and edited strains. B) *APT*-sgRNA-g1 and g2 were cloned individually and together into the PHYCUT plasmid and conjugated into *P. tricornutum*. APT knock outs were selected for on 50% salt L1 solid media with 10 $\mu$M 2-FA. The guides and screening primers used are indicated by the diagram above each set of agarose gels of PCR products (top) and T7E1 assays (bottom). Sizes (in bp) are indicated next to the gel images. Black horizontal lines separate colonies derived from two independent conjugations. WT indicates PCR amplicons and T7E1 digests from untreated *P. tricornutum*. T7E1 digests contain a mixture of PCR amplicons from WT and edited strains. C) The knock out efficiency of *APT* in *P. tricornutum* exconjugants as determined by the ratio of exconjugants on solid media containing 2-FA or zeocin. Barplots are mean of 3 biological replicates with error bars representing standard error of the mean (* p ¡ 0.05). D) Sequences of edited *APT* genes from the indicated exconjugants. The dashed light grey line indicates the spacial separation between the two target sites. Total size of deletion is indicated in parentheses for deletions extending outside of the region shown.

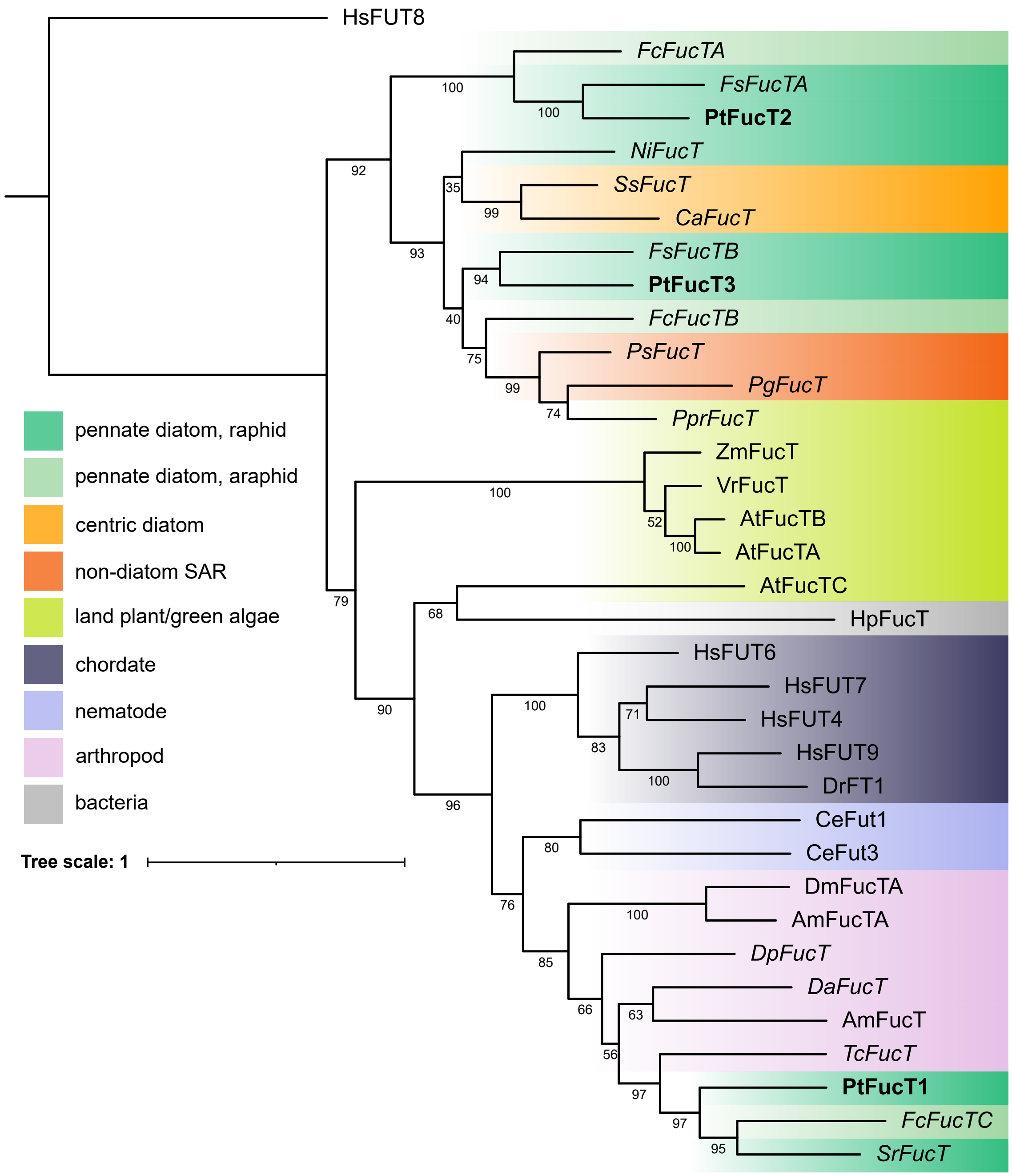


**Figure S 3.** Phylogenetic analysis of the three GT10-family PtFucTs. The globular domains of GT10-family FucTs were aligned with MAFFT and assembled into a phylogenetic tree with IQtree using 100 bootstrap replicates, bootstrap values are shown for each branch point. HsFUT8, a GT37-family FucT, is included as an outgroup. The three PtFucTs are shown in bold. FucTs in non-italicized font are characterized in the CAZy database, FucTs in italics are un-characterized homologs identified in this study.

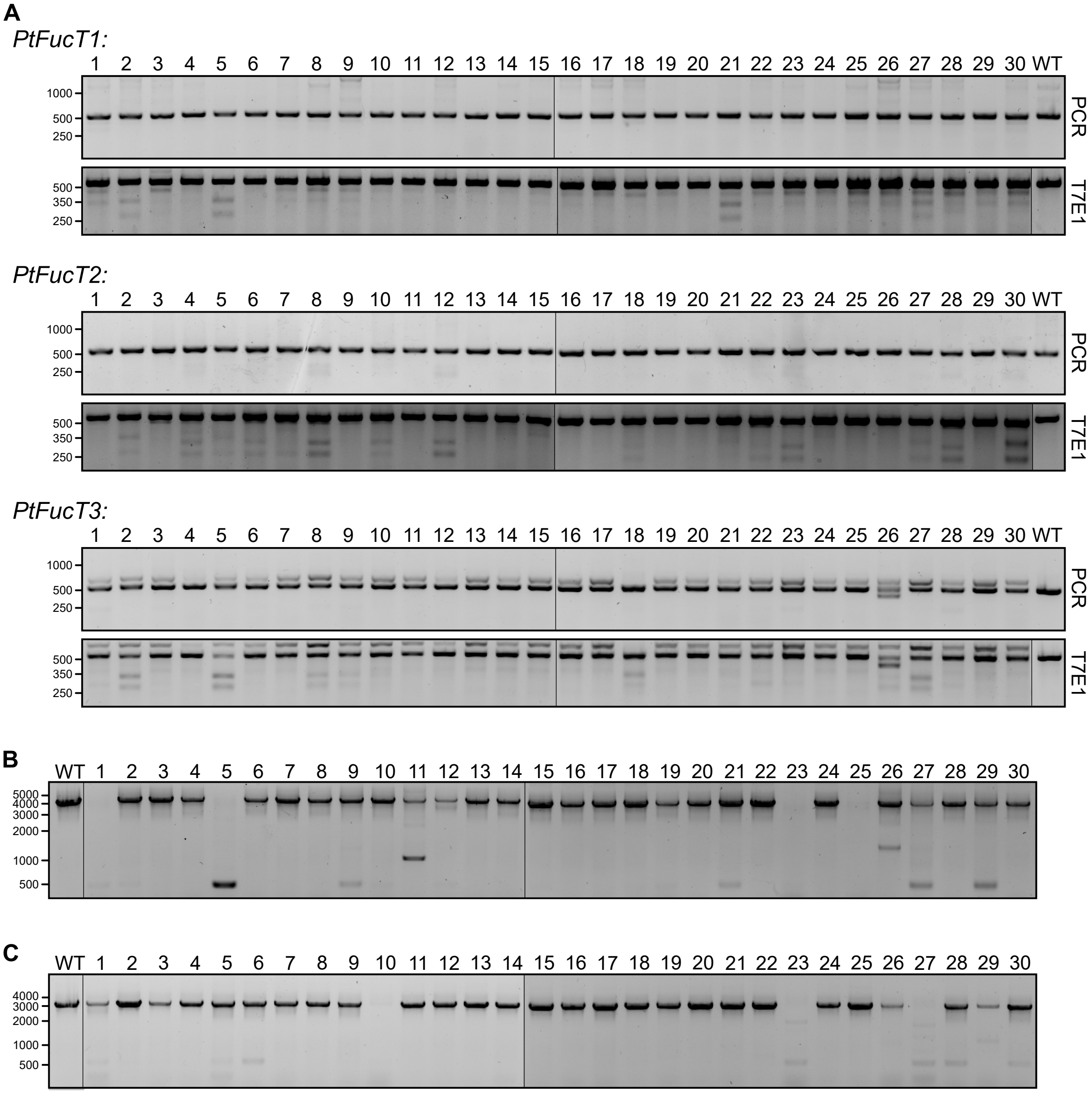


**Figure S 4.** FucT PHYCUT exconjugants screened for the presence of editing at targeted loci. A) Short-range PCRs and T7 Endonuclease (T7E1) assays of the three target genes on for all 30 exconjugants screened. Sizes (in bp) are indicated next to the gel images. WT indicates PCR amplicons and T7E1 digests from untreated *P. tricornutum*. T7E1 digests contain a mixture of PCR amplicons from WT and edited strains. B) Long-range PCR surrounding target sites for *PtFucT2* and *3* show large deletions between the target sites. The amplicon size of a deletion from the *PtFucT2*-sgRNA-tg to *PtFucT3*-sgRNA-tg target sites is 414bp. C) Long-range PCR surrounding the *PtFucT1*-sgRNA-tg target site revealed presence of larger deletions.

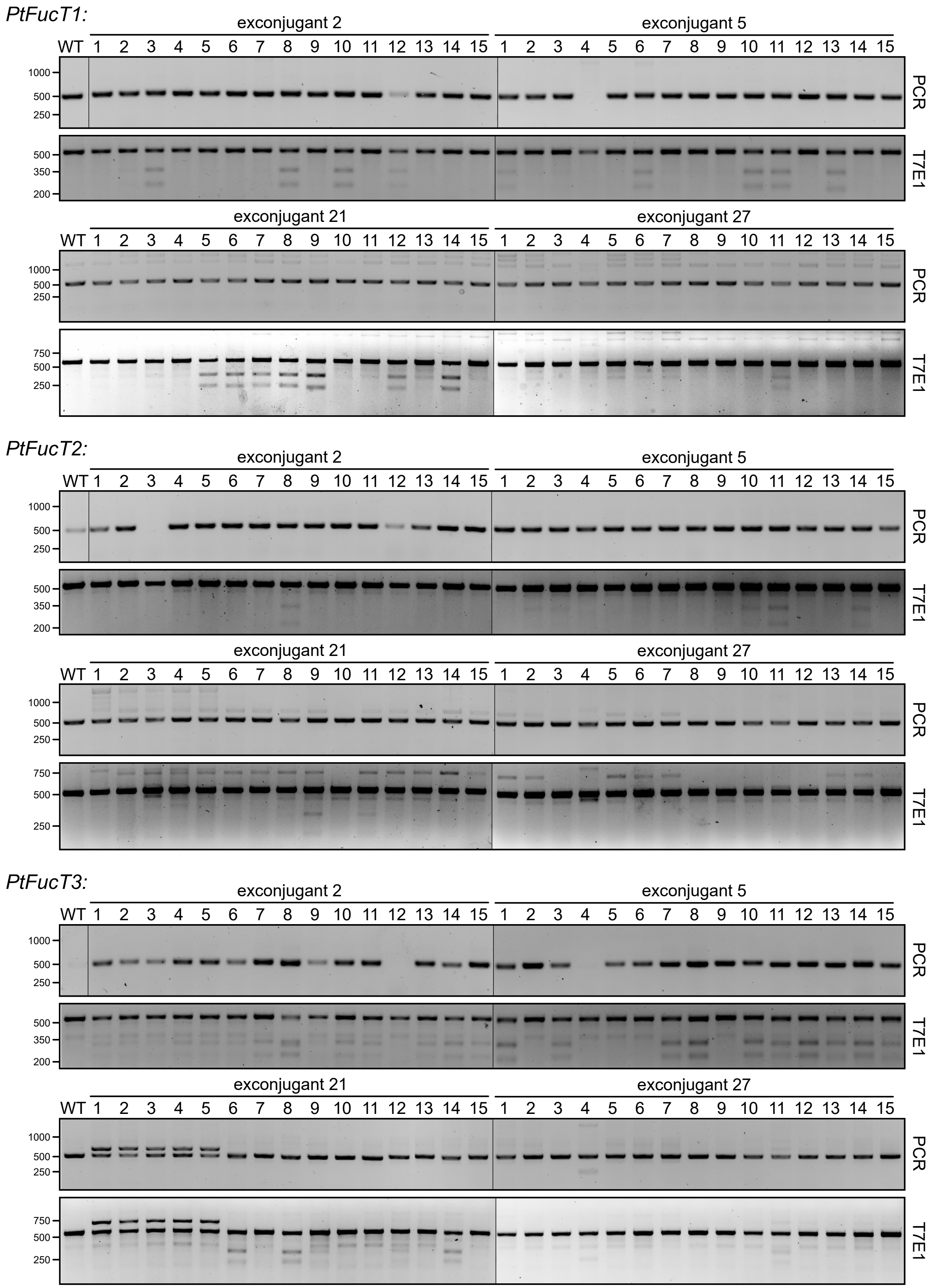


**Figure S 5.** FucT PHYCUT subclones screened for the presence of editing at targeted loci. Short-range PCRs and T7 Endonuclease (T7E1) assays of the three target genes for 15 subclones each derived from the exconjugants 2, 5, 21, and 27. Sizes (in bp) are indicated next to the gel images. WT indicates PCR amplicons and T7E1 digests from untreated *P. tricornutum*. T7E1 digests contain a mixture of PCR amplicons from WT and edited strains.

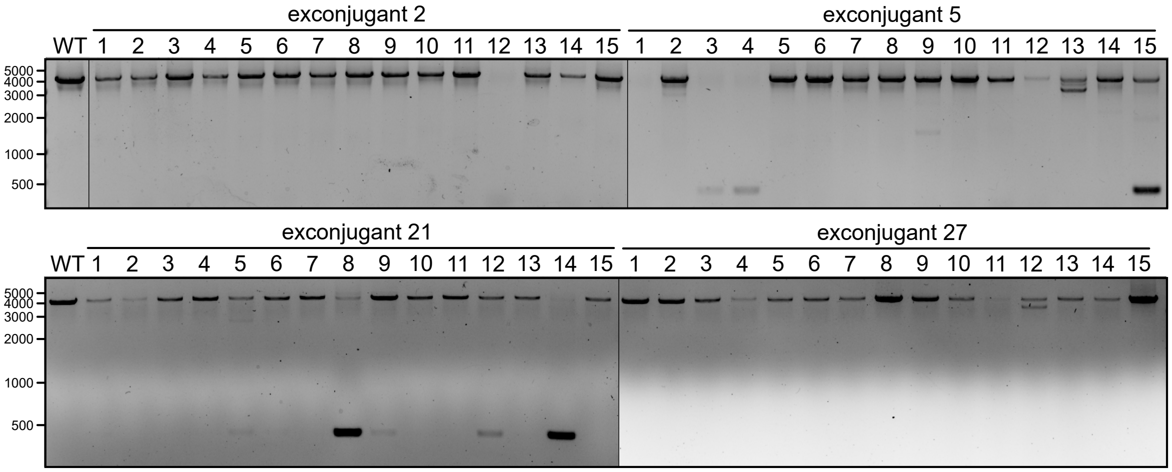


**Figure S 6.** FucT PHYCUT subclones screened for defined-length deletions between adjacent target sites. 15 subclones each derived from the exconjugants 2, 5, 21, and 27 are screened. Long-range PCR surrounding target sites for *PtFucT2* and *3* show large deletions between the target sites. Sizes (in bp) are indicated next to the gel images. WT indicates PCR amplicons from untreated *P. tricornutum*. The amplicon size of a deletion from the *PtFucT2*-sgRNA-tg to *PtFucT3*-sgRNA-tg target sites is 414bp.

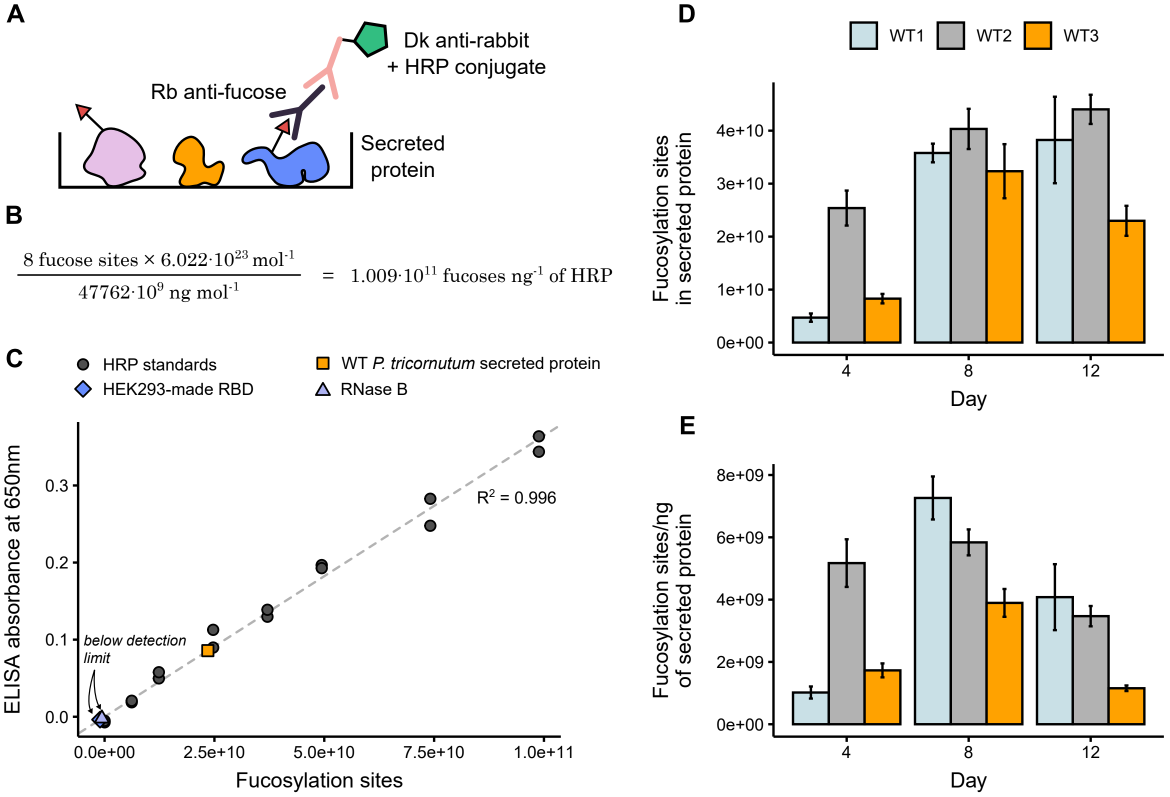


**Figure S 7.** Indirect ELISA protocol allows sensitive levels of fucosylation detection. A) Protein samples from either extracellular media of *P. tricornutum* or purified control samples were passively adhered to the assay plates, where $\alpha$(1,3)-fucose epitopes were detected by a rabbit (Rb) mAb. A secondary donkey (Dk) anti-rabbit HRP-conjugated Ab was then used to amplify and emit a colourimetric signal. B) The glycoprotein horseradish peroxidase (HRP) was used to create standard curves for all ELISA experiments. HRP contains 8 $\alpha$(1,3)-fucosylated N-glycosites, which was used in conjunction with the average molecular weight of the HRP glycoprotein to calculate the fucose sites per ng of HRP. C) An example standard curve using standards from 0-20 ng/mL of HRP shows an excellent linear relationship, with wild type *P. tricornutum* secreted protein samples falling within this range. The controls of 50 ng/$\mu$L HEK293-made RBD (human complex glycans containing $\alpha$(1,6)-core fucosylation), and 50 ng/$\mu$L RNase B (oligomannose glycans) were included, which both showed zero detection in the assay. D) Fucosylation levels of the three wildtype *P. tricornutum* samples included in this study on days 4, 8, and 12. Barplots are mean of 3 biological replicates with error bars representing standard error of the mean. E) Fucosylation levels of the three wildtype P. tricornutum samples included in this study, normalized by protein concentration in the extracellular media, on days 4, 8, and 12. Barplots are mean of 3 biological replicates with error bars representing standard error of the mean.

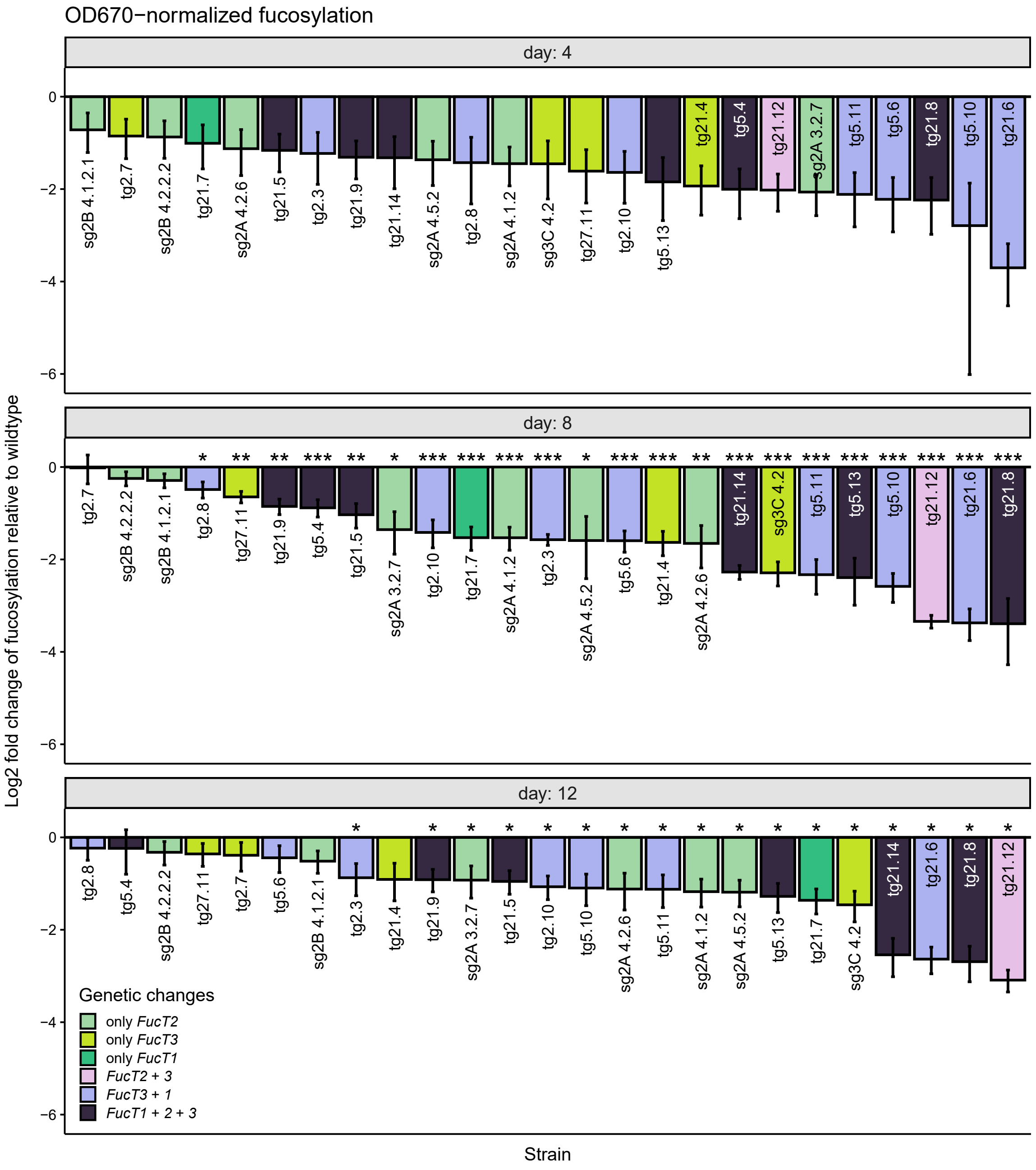


**Figure S 8.** Culture density-normalized fucosylation of secreted proteins relative to wild type. $\alpha$(1,3)-fucosylation of glycoproteins secreted into the extracellular media by FucT mutant strains, relative to wild type *P. tricornutum*, plotted as log2 fold change. Degree of fucosylation on days 4, 8, and 12 of growth was determined via indirect ELISA for the core $\alpha$(1,3)-fucose motif and normalized to culture density as determined by absorbance at 670 nm. Strains are coloured by their genetic edits. Barplots are means of 3 biological replicates relative to 9 biological replicates of wildtype *P. tricornutum* with error bars representing standard error of the mean. P-values compared to the wild type average for each day are displayed above each sample (* $p < 0.05$, ** $p < 0.01$, *** $p < 0.001$)

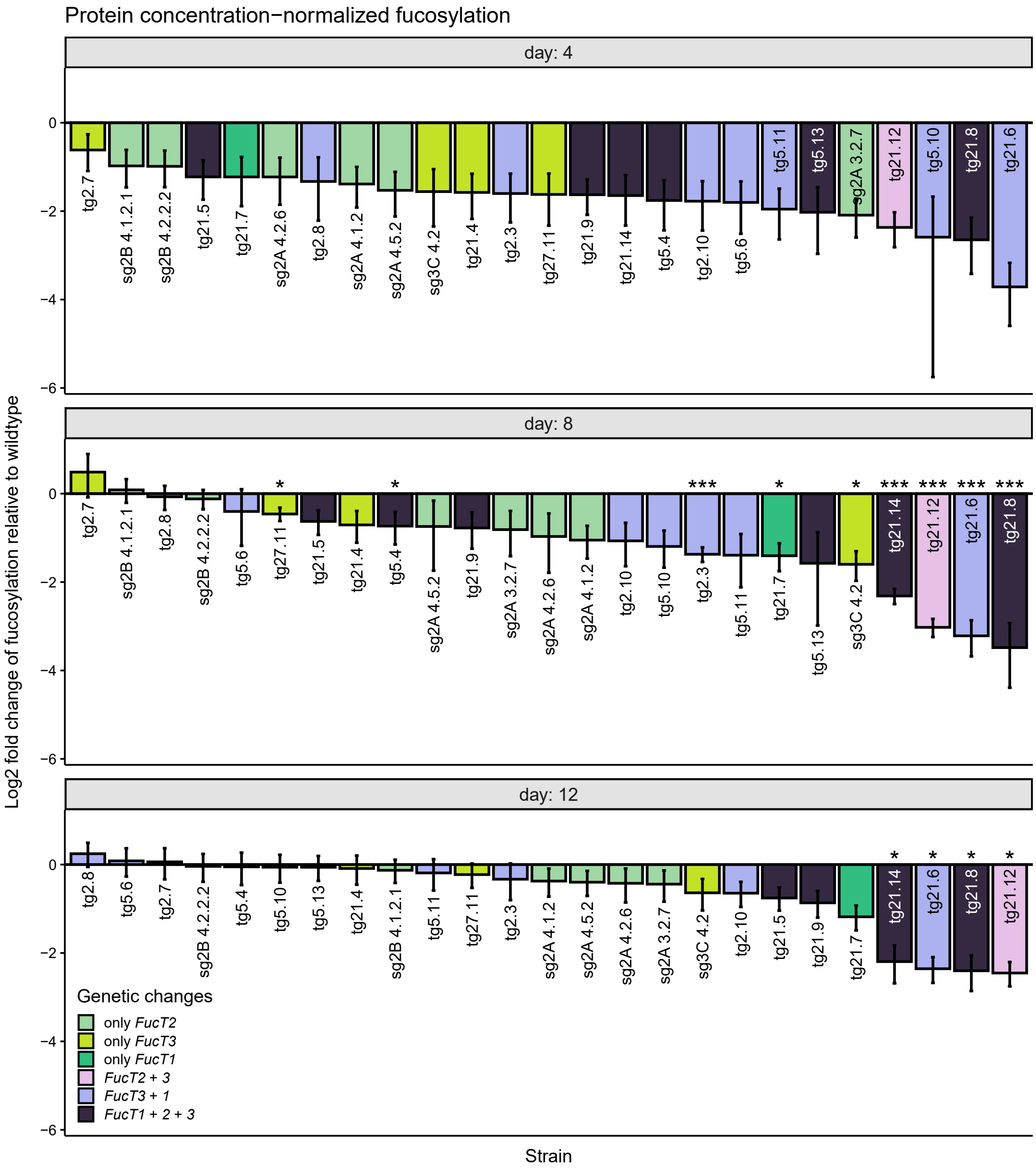


**Figure S 9.** Protein-normalized fucosylation of secreted proteins relative to wild type. $\alpha$(1,3)-fucosylation of glycoproteins secreted into the extracellular media by FucT mutant strains, relative to wild type *P. tricornutum*, plotted as log2 fold change. Degree of fucosylation on days 4, 8, and 12 of growth was determined via indirect ELISA for the core $\alpha$(1,3)-fucose motif and normalized to cprotein concentration in the extracellular media as determined by Bradford assay. Strains are coloured by their genetic edits. Barplots are means of 3 biological replicates relative to 9 biological replicates of wildtype *P. tricornutum* with error bars representing standard error of the mean. P-values compared to the wild type average for each day are displayed above each sample (* $p < 0.05$, ** $p < 0.01$, *** $p < 0.001$)

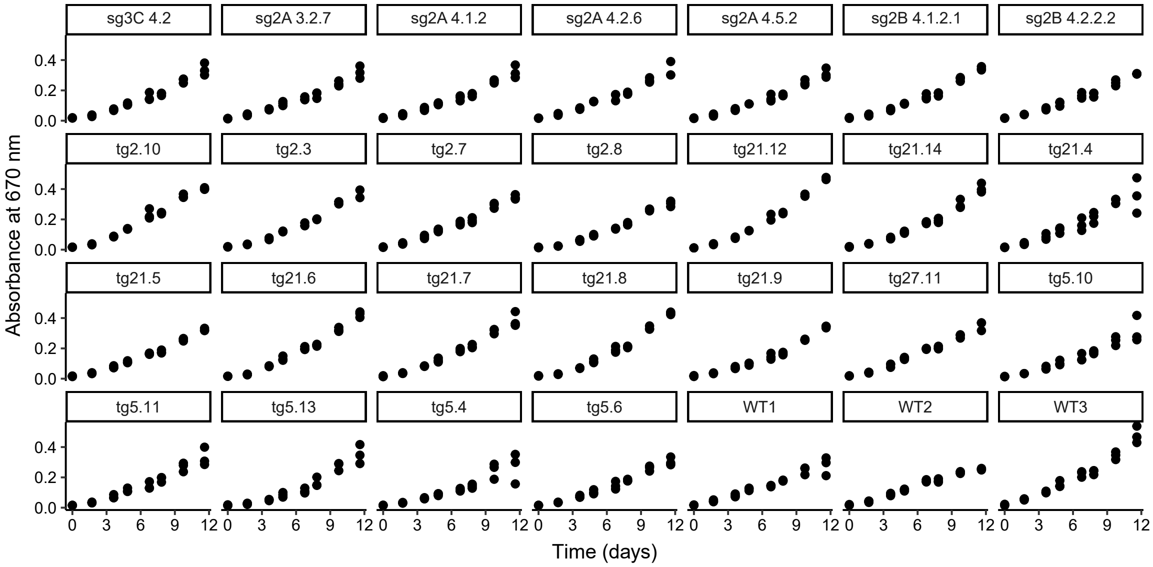


**Figure S 10.** Growth curves for all strains studied in ELISA experiment. Three replicates for each strain were grown throughout the 12-day time course with the absorbance of the culture at 670 nm measured every other day. Each point represents the average for each biological replicate.
